# Molecular signatures of cortical expansion in the human fetal brain

**DOI:** 10.1101/2024.02.13.580198

**Authors:** G. Ball, S. Oldham, V. Kyriakopoulou, L. Z. J. Williams, V. Karolis, A. Price, J. Hutter, M.L. Seal, A. Alexander-Bloch, J.V. Hajnal, A.D. Edwards, E.C. Robinson, J. Seidlitz

## Abstract

The third trimester of human gestation is characterised by rapid increases in brain volume and cortical surface area. A growing catalogue of cells in the prenatal brain has revealed remarkable molecular diversity across cortical areas.^1,2^ Despite this, little is known about how this translates into the patterns of differential cortical expansion observed in humans during the latter stages of gestation. Here we present a new resource, μBrain, to facilitate knowledge translation between molecular and anatomical descriptions of the prenatal developing brain. Built using generative artificial intelligence, μBrain is a three-dimensional cellular-resolution digital atlas combining publicly-available serial sections of the postmortem human brain at 21 weeks gestation^3^ with bulk tissue microarray data, sampled across 29 cortical regions and 5 transient tissue zones.^4^ Using μBrain, we evaluate the molecular signatures of preferentially-expanded cortical regions during human gestation, quantified *in utero* using magnetic resonance imaging (MRI). We find that differences in the rates of expansion across cortical areas during gestation respect anatomical and evolutionary boundaries between cortical types^5^ and are founded upon extended periods of upper-layer cortical neuron migration that continue beyond mid-gestation. We identify a set of genes that are upregulated from mid-gestation and highly expressed in rapidly expanding neocortex, which are implicated in genetic disorders with cognitive sequelae. Our findings demonstrate a spatial coupling between areal differences in the timing of neurogenesis and rates of expansion across the neocortical sheet during the prenatal epoch. The μBrain atlas is available from: https://garedaba.github.io/micro-brain/ and provides a new tool to comprehensively map early brain development across domains, model systems and resolution scales.

---

The human cortex is a tapestry of specialised cortical areas supporting diverse and complex behaviours, each identifiable on the basis of distinct patterns of cyto-architecture, chemo-architecture, and axonal connectivity.^6–10^ During gestation, waves of neurons are generated from progenitor cells lining the cerebral ventricles and migrate outwards along supporting radial glia to form the layers of the cortex.^11–13^ Prior to the ingress of extrinsic connections via the thalamus,^14^ the progressive differentiation of cortical areas is orchestrated by transcription factors expressed along concentration gradients and translated from the ventricular zone (VZ) to secondary progenitors of the subventricular zone (SVZ), then onto neurons in the cortical plate (CP), forming functional territories.^2,11,15–18^ This process follows a precise spatiotemporal schema,^11,18–22^ the traces of which extend far beyond the nascent stages of neurogenesis and are echoed in patterns of cytoarchitecture, axonal connectivity and function.^23–29^

Focused on uncovering the mechanisms that govern areal differentiation, studies have begun to catalogue the cellular diversity of the developing human cortex, and genes that encode it, with increasing granularity and scale.^1,2,30–32^ Regional specialisation of cell types has been observed from early in gestation, with diversity of cortical gene transcription most evident in mid- to late-gestation but persisting into adulthood and aligning with structural and functional organisation of the brain.^4,26,33–36^

The third trimester of human gestation is characterised by rapid and sustained increases in brain volume and cortical surface area.^11,37,38^ Differential rates of areal expansion during human development mirror evolutionary trends in cortical scaling and function^39–44^ with preferential expansion in areas vulnerable to disruption in neurodevelopmental,^45^ neurological^46^, genetic^47^ and psychiatric^48^ disorders. Juxtaposed hypotheses implicate either the production of glia^49,50^, or neurons^51–54^ from specialised progenitor populations of the outer SVZ, in the expansion of the primate cortex. Thus, the distribution of distinct cell populations across the developing cortex may mediate areal differences in expansion and vulnerability to insult^18,33^ but we currently do not have a clear understanding of how this molecular diversity is translated into cortical organisation in humans *in vivo*.

## μBrain: A three-dimensional microscale atlas of the fetal brain

To bridge this gap, we sought to construct a 3D digital atlas of the developing brain at micrometre scale using a public resource of 81 serial histological 2D sections of a prenatal human brain at 21 postconceptional weeks (PCW).^3,4^ Source data included serial coronal sections (20μm thickness) obtained from the right hemisphere of a single prenatal brain specimen (21 PCW; female), Nissl-stained, imaged at 1 micron resolution and labelled with detailed anatomical annotations, alongside interleaved coronal sections stained with *in situ* hybridisation (ISH) of n=41 developmental gene markers, as reported by Ding et al^3^ (**Figure S1**; **Table S1-S3;** see **Methods**). In this and 3 other specimens (15, 16 and 21 PCW, 2 female), anatomical annotations had been used to guide a series of laser microdissections (LMD) across multiple cortical areas and layers of the cortical anlage (e.g.: cortical plate, subplate, intermediate zone, ventricular zone; **Table S4**) in the left hemisphere to measure regional gene expression via RNA microarrays, as described by Miller et al.^4^ Nissl- and ISH-stained sections with corresponding anatomical labels and LMD arrays were made available as part of the BrainSpan Developing Brain Atlas [https://atlas.brain-map.org/atlas?atlas=3].

Artefacts due to tissue preparation, sectioning and staining procedures (including tearing and folding of sections) are common in histological data and can present difficulties for downstream processing pipelines.^55–58^ To correct for tissue artefacts present in the histological data, we designed an automated detect-and-repair pipeline for Nissl-stained sections based on *pix2pix*, a Generative Adversarial Network (GAN)^59,60^ (**Figure 1a-d; Supplemental Methods**). Using 256 × 256 pixel image patches drawn from 73/81 labeled histological sections (n=8 reserved for model testing) with paired anatomical labels, we trained a GAN model to produce Nissl-contrast images conditioned on a set of 20 tissue labels (**Figure 1a; Table S2**). After training, the model was able to produce realistic, Nissl-stained image patches matched on colour hue and saturation to the original data using tissue annotations alone (**Figure 1c**). Model performance was robust to different parameter settings and model architectures (**Figure S2**). Using the trained model, we generated synthetic Nissl-contrast image predictions from anatomical annotations of each section and identified artefacts in the histological data based on deviations in pixel hue and saturation from the model prediction. Outlier pixels were replaced with model predictions using Poisson image editing^61^ (**Figure 1d**) resulting in n=79 (2 excluded due to extensive missing tissue) complete histological sections (**Figure S1; Table S1**).

**Figure 1:**
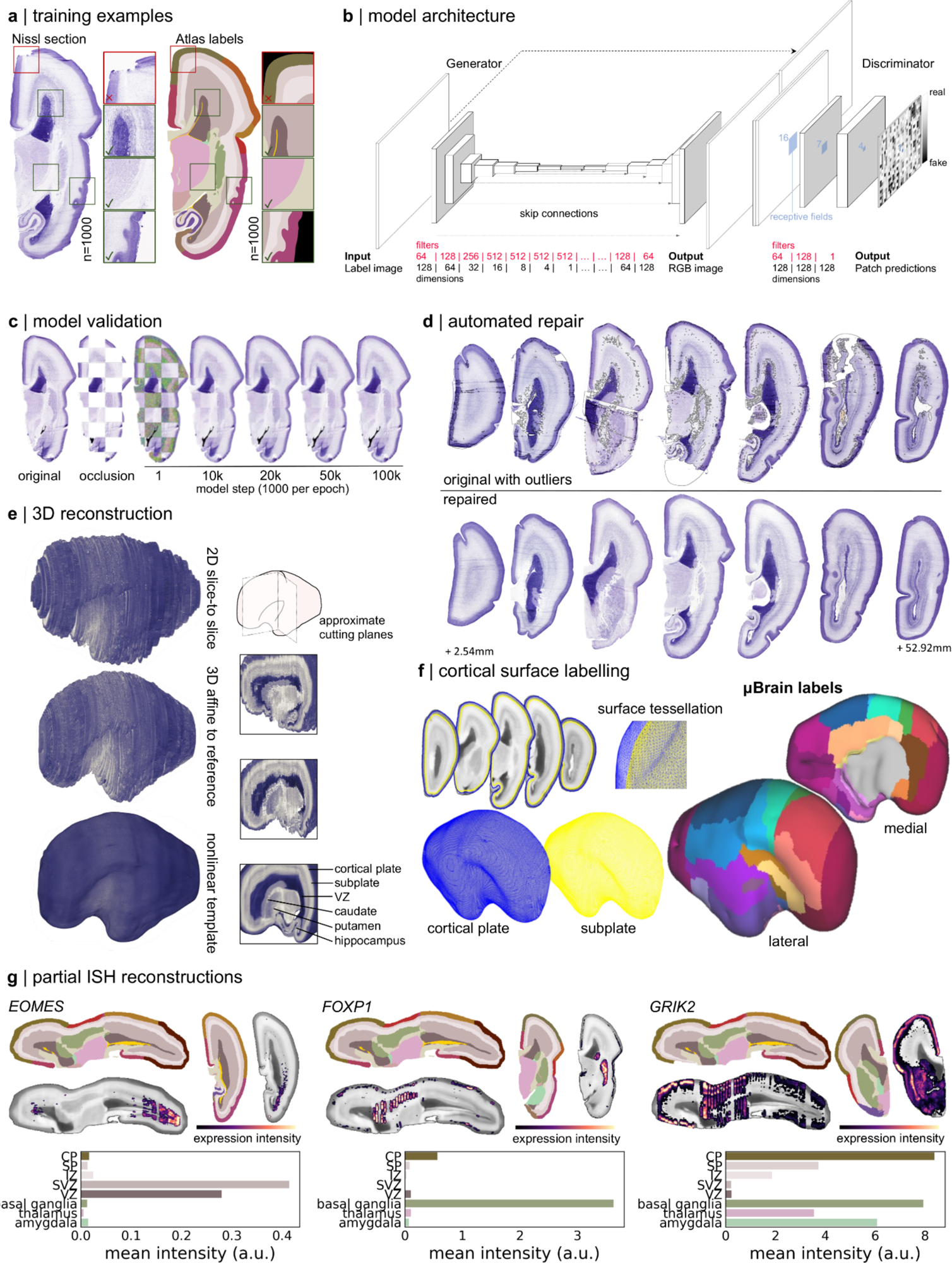
Generation of a 3D anatomical atlas of the mid-gestation fetal brain. **a**. Paired histological sections and simplified anatomical annotations were divided into 256 × 256 random patches (n=1000) for model training. Patches were quality checked prior to selection to ensure good overlap between labels and anatomy and no tissue damage. **b**. *Pix2pix* model architecture showing a U-Net generator coupled with a PatchGAN discriminator. Box sizes represent image width, height and number of filters/channels (depth) at each layer. Filters and dimensions of each layer are shown below. **c**. Model performance was evaluated on a set of sections that were not included in the training dataset. Checkerboard occlusions are shown with the original section, occluded patch predictions are shown using the trained model after a given number of iterations. **d**. The trained model was used to replace RGB values of outlying pixels with synthetic estimates. Top row: original sections spaced throughout the cerebral hemisphere with automatically identified outlier pixels outlined in grey. Bottom row, repaired sections. **e**. Repaired sections were aligned via linear, affine and iterative nonlinear registrations (see **Methods**) to create a 3D volume with final isotropic resolution of 150um. Right: Cut-planes illustrate internal structures after each stage of reconstruction. The reconstructed tissue label volume is shown in **Figure S4**. **f**. The outer (pial) and inner (subplate) cortical plate boundaries were extracted as surface tessellations. The μBrain cortical labels were projected onto the surface vertices to form the final cortical atlas (see **Figure S4**). Cortical areas correspond to matched LMD microarray data (**Table S4**; **Figure S4d**). **g**. Partial reconstructions of *EOMES*, *FOXP1* and *GRIK2* ISH data. ISH stained sections were registered to nearest Nissl-stained sections and aligned to the μBrain volume. Top row: selected axial and coronal sections of the μBrain volume and corresponding tissue labels with ISH expression of three developmental genes: *EOMES, FOXP1* and *GRIK2* overlaid. Expression intensity was derived from false-colour, semi-quantitative maps of gene expression. Bottom row: average expression intensity within each tissue or brain structure based on μBrain tissue labels. Averages were calculated only within sections where ISH was available for each gene.

Histological atlases of the cerebral cortex^6,10^ have proven invaluable for understanding human brain organisation but are limited by the loss of spatial information inherent to 2D representations of 3D structures. Reconstructions of 3D brain volumes from serial tissue sections of post mortem tissue allow the examination of intact brain anatomy at a scale inaccessible to current neuroimaging technologies.^62^ We combined repaired tissue sections into a 3D volume of the right hemisphere using iterative affine image registration constrained by a tissue shape reference derived from fetal MRI (**Figure S3; Supplemental Methods**),^63^ followed by nonlinear alignment to account for warping between adjacent sections. Using the aligned data, we generated a 3D volume resampled to voxel resolution 150 × 150 × 150*µ*m with dimension 189 × 424 × 483 voxels (28.35 × 63.60 × 72.45mm) (μBrain; see **Methods**; **Figure 1e, Figure S4a-b)**. Following reconstruction, we benchmarked the size of the reconstructed μBrain volume against standard fetal growth metrics for a 23 week (gestational age; GA, equivalent to 21 PCW) fetus (μBrain length = 62.7mm, 23 week GA occipital-frontal diameter median [5^th^, 95^th^ centile] = 73.3 mm [68.2, 78.5]),^64^ and compared tissue volume estimates based on reconstructed anatomical labels (parenchymal volume = 25.8ml, right hemisphere) to previously reported 3D MRI-derived fetal brain volumes (supratentorial volume [both hemispheres] at 23 week GA = 60.26ml).^65^ Adapting protocols from neuroimaging analysis, we extracted the inner and outer surfaces of the cortical plate and projected a set of 29 cortical area labels derived from the histological tissue annotations (**Table S2**) onto the surface vertices to form the μBrain cortical atlas (**Figure 1f**). The μBrain cortical atlas represents a new parcellation of the developing brain defined according to the hierarchical ontology of the reference annotations and matched to corresponding LMD microarray data (**Table S2, S4; Figure 4c-d**).

In addition to the whole brain volume and cortical atlas, we created partial 3D reconstructions of ISH staining for 41 genes (see **Methods**; **Figure 1g**). Based on an average 41 tissue sections per gene (**Table S3**), semi-quantitative maps of gene expression revealed the tissue- and region-specific distributions of several genes, including caudal enrichment of the transcription factor *EOMES* in the subventricular zone,^15^ and markers of neuronal migration (*DCX*^66^) and synaptic transmission (*GRIK2*^67^), in the cortical plate (**Figure 1g**; **Figure S5**)

Existing histological brain atlases, including those of the adult human,^62,68,69^ mouse,^70,71^ and macaque^72^ brains facilitate integration with other data modalities, including neuroimaging, and are amenable to advanced computational image analysis methods to extract quantitative measures of neuroanatomy across multiple scales.^73,74^ Building upon existing resources,^3,4^ we have created the μBrain atlas (**Figure 1; Figure S4**), a new and freely-available 3D volumetric model of the 21 PCW fetal brain at 150*µ*m resolution, accompanied by a set of n=20 cerebral tissue labels (**Figure S4a-b**); surface models of the cortical plate surface and cortical plate/subplate interface with n=29 cortical area labels (**Figure 4c**) and n=41 partial reconstructions of ISH expression data (**Figure 5**). Cortical areas are matched to normalised gene expression data from corresponding LMD microarrays (**Table S4; Figure S4d**) across multiple tissue zones in three additional prenatal specimens (total n=4) providing a 3D anatomical coordinate space to facilitate integrated imaging-transcriptomic analyses of the developing brain. Below, we use the μBrain atlas to evaluate the molecular and cellular correlates of cortical expansion in the third trimester of human gestation.

## Tissue- and region-specific gene expression in the mid-gestation brain

We sought to characterise patterns of gene expression in the mid-gestation brain and identify developmental and region-specific genes with putative roles in cortical expansion. To do so, we used publicly-available microarray data from four prenatal brain specimens aged 15 to 21 PCW.^4^ Microarray probe annotations were updated and tissue samples matched to the μBrain atlas (**Table S4**) yielding expression data of 8771 genes sampled from between 18 and 27 brain regions and across 5 tissue zones for each specimen (see **Methods**; **Figure S4d**). Applying PCA to gene expression profiles, we found that tissue samples were primarily separated according to location in mitotic (VZ, SVZ) or post-mitotic tissue zones, rather than across regions (**Figure 2a**)^4^ – a pattern that was replicated across all specimens when analysed separately (**Figure S6**). Focusing on expression profiles within each tissue zone, samples clustered according to maturity (**Figure 2a**; **Figure S7**) with developmental changes in gene expression most similar across adjacent mitotic (CP and SP, r= 0.67) and post-mitotic zones (SVZ and VZ, r=0.43; **Figure S8**). In line with evidence of a transition in VZ cell fate around mid-gestation,^75^ we observed increased expression of genes enriched in post-mitotic excitatory neurons and interneurons (e.g.: *GRIK1-3*; *GLRA2*; *SCN3B*) between 15 and 21 PCW.^1^ In the SP and CP, this transition coincided with an increase in genes expressed by radial glia (*BMP7*; *SOX3*) and oligodendrocyte precursor cells (OPCs; *CA10*) with a transitory decrease in microglia-enriched genes in the CP (*GPR34*; *TREM2*)^76^ (**Figure 2b; Table S5**).

**Figure 2:**
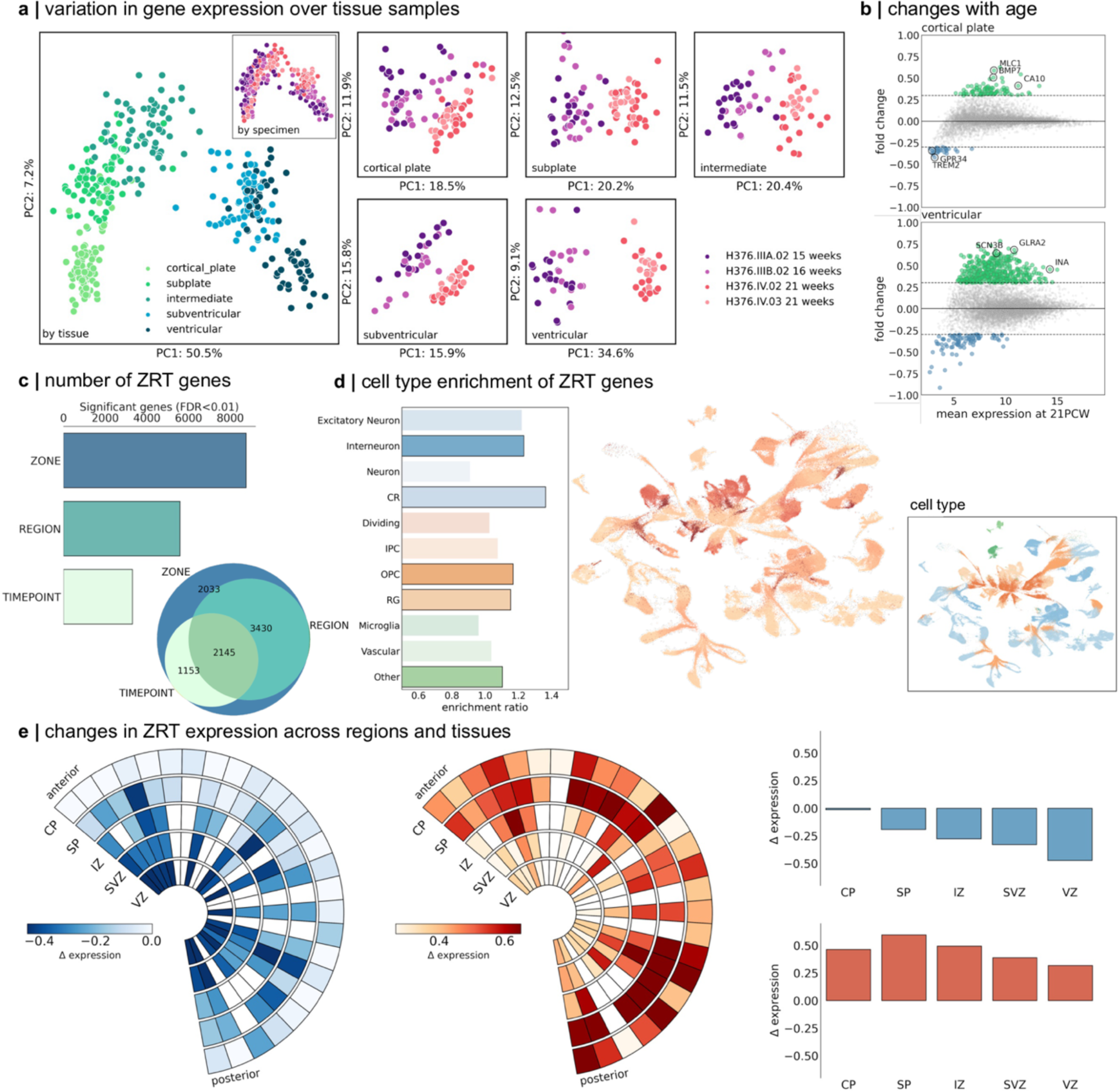
Regional gene expression in the mid-gestation fetal brain. **a.** PCA of LMD microarray data (n=8771 genes) in four prenatal brain specimens aged 15 PCW to 21 PCW. All tissue samples are shown (left) coloured by tissue zones (main) and specimen (inset). PCA was applied to all samples in each tissue zone separately (right). Samples are coloured by specimen and cluster by age. **b**. PC1 was associated with age-related change in all tissues and correlated between neighbouring zones. Plots show mean gene expression at 21 PCW (averaged over specimen and region) against fold change in gene expression between 15/16 PCW and 21 PCW for two tissue zones (cortical plate, top and ventricular zone, bottom). Genes with a log2(fold change) > 0.3 are shown in green (<-0.3 in blue). Representative genes are highlighted. **c.** Number of genes with differential expression over tissue zones (ZONE), cortical region (REGION) or timepoint (TIME). Venn diagram shows overlap of gene sets. In total, n=2145 were differentially expressed across zone, region and time (ZRT genes). **d.** enrichment of ZRT genes in cell types previously identified in the mid-gestation fetal brain (left).^2^ UMAP projection of cell types showing enriched clusters of OPCs and radial glia. Inset: UMAP projection coloured by cell type. **e.** ZRT gene expression over time and region. Wedge plots (left) show the pattern of expression of ZRT genes that decrease (left) or increase (right) between 15 and 21 PCW. Rows indicate tissue zones and columns indicate cortical regions ordered from anterior to posterior poles. Boxes are coloured by change in gene expression over time (*Δ* expression). Right: bar charts show mean change in gene expression for decreasing (top) and increasing (bottom) ZRT genes averaged within tissue zones.

Across all tissue samples, we tested for differences in gene expression across zones (CP, SP, etc.), regions (motor, sensory, etc) and timepoints (early vs mid-gestation). This resulted in a subset of n=2145 (24.5%) genes with differential expression across all three factors, termed Zone-Region-Tissue (ZRT) genes (p<0.01 after FDR correction; **Figure 2c; Table S6)**. We reasoned that this subset, characterised by genes with dynamic regional and temporal expression in mid-gestation, would be associated with differential rates of cortical expansion during development. To support this line of reasoning, we found that the ZRT cluster was enriched for genes upregulated in the third trimester^77^ (enrichment ratio = 1.89, hypergeometric test p_hypergeom_<0.0001) and highly expressed in adolescent and adult brain tissue, compared to non-ZRT genes (**Figure S9**). ZRT genes included several human transcription factors (e.g.: *EGR1*, *JUNB*, *ZNF536*)^78^ and were significantly enriched in radial glia (*SOX2*, *HES5,* p_hypergeom_=0.03176), OPCs (*OLIG1*, *PDGFRA;* p_hypergeom_=0.0424) and migrating interneurons (*CALB2*, *CNR1*; p_hypergeom_=0.0009; **Figure 2d; Table S7**).^2^ We observed highest ZRT expression in the subplate, with increasing expression of ZRT genes in postmitotic zones (CP, SP and IZ) compared to the SVZ and VZ, between 15 and 21 PCW (**Figure 2e**). Examining ZRT gene annotations revealed enrichment of critical neurodevelopmental functions including cell-cell adhesion (GO: 0098742; *CHD1*, *EFNA5*, *NLGN1*, *NRXN1;* p_FDR_<0.0001, background set = 8771 genes), forebrain development (GO: 0030900; *CASP3*, *CNTN2*, *DLX2*, *FOXP2*, *NEUROD6*; p_FDR_=0.026) and neuron projection guidance (GO: 0097485; *EFNA2*, *EFNA5*; p_FDR_ = 0.0034) (**Table S8**). The ZRT geneset was additionally enriched for high-confidence ASD-linked genes (n = 43, p_hypergeom_ = 0.034)^79^ including *SCL6A1*, *CACNA1C* and *CHD7* and pathogenic variants in 161 ZRT (7.5%) genes have been linked to neurodevelopmental and cognitive phenotypes and brain malformations^80^ including *MAGEL2* (Schaaf-Yang syndrome^81^), *AFF2* (Fragile-X-E^82^) and *ADGRG1* (polymicrogyria^83^) (**Table S9**)

## Regional differences in the rate of cortical expansion *in utero* during the third trimester

We hypothesised that the dynamic temporal and regional patterning of ZRT genes across tissue zones support differential rates of areal expansion across the cortex. To test this, we acquired n=240 motion-corrected fetal brain MRI scans from 229 fetuses aged between 21^+1^ and 38^+2^ gestational weeks^+days^ as part of the Developing Human Connectome Project (dHCP).^84^ Volumetric T2-weighted scans were automatically reconstructed to 0.5mm isotropic voxel resolution^85,86^; then tissue segmentations were initially extracted using neonatal protocols^87^, followed by extensive manual editing to ensure accuracy (**Methods**). Manually-corrected segmentations were then used to generate cortical surface reconstructions (**Figure 3a**).^88^ For analysis, individual cortical surfaces were aligned to a fetal spatiotemporal atlas using a nonlinear, biomechanically-constrained surface registration (Multimodal Surface Matching [MSM]; **Figure 3a-c**).^89–92^ At each stage, outputs were visually quality-checked and any failures removed. In total, data from n=195 scans in 190 fetuses (gestational age: 21^+1^ - 38^+2^ weeks; 88 female) were included in the analysis (**Figure S10**).

**Figure 3:**
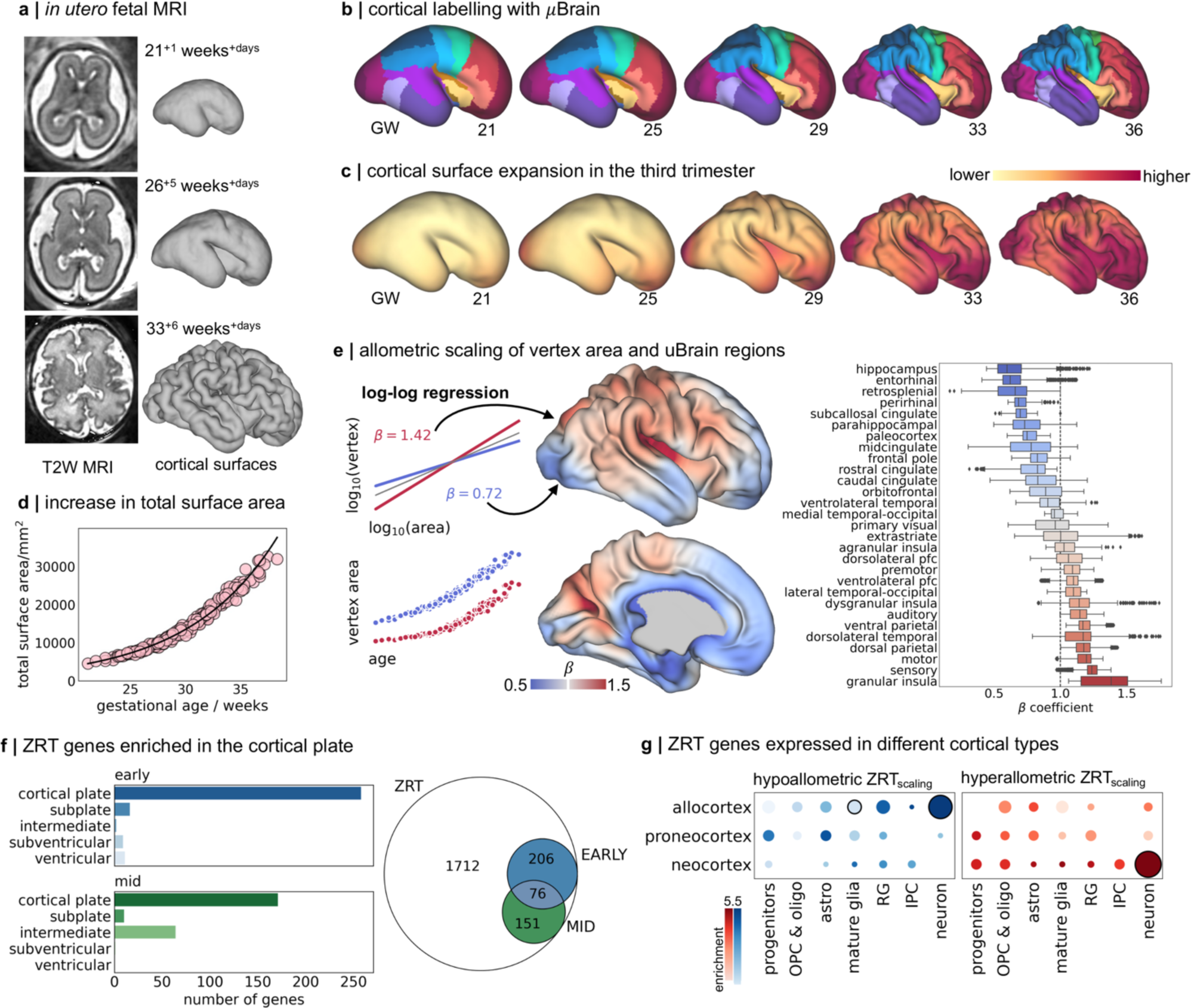
Preferential cortical expansion during the third trimester. **a.** n=195 fetal MRI scans were acquired during the third trimester of pregnancy. T2-weighted (T2W) scans were reconstructed using a motion-robust processing pipeline and used to generate tessellated cortical surface representations that were aligned to the dHCP fetal surface template **b**. μBrain cortical labels projected onto dHCP fetal template surfaces from 21 to 36 weeks gestation using nonlinear surface registration. Surfaces are scaled to the same size for visualisation. **c.** For each timepoint, weighted average vertex area maps are displayed on the respective surface templates. Fetal cortical area maps were calculated from individual, co-registered and resampled fetal surfaces using a Gaussian kernel (sigma = 1 week). **d**. total cortical surface area calculated across all surface vertices (excluding the midline) as a function of gestational age at scan. **e.** Left: Models of allometric scaling were calculated for each vertex, modelling log10(vertex area) as a function of log10(total area)(top). In this framework, *β*>1 indicates hyperallometric growth (a relative expansion faster than the global rate). Note that a faster growth rate does not necessarily equate to greater total area at any given time (bottom). Middle: Hyperallometric scaling with respect to total cortical surface area (***β*** > **1**) plotted on the 36w template surface representing preferential cortical expansion during development. Right: Distribution of scaling coefficients for all vertices in each μBrain label in **a**, ordered by mean scaling. **f.** Right: In total, expression of 433 ZRT genes were correlated with areal scaling in gestation. Left: Significant associations (Kendall’s *τ*, pFDR<0.01) were observed across both early (15/16 PCW, n=2) and mid-gestation (21 PCW, n=2) timepoints and in all tissue zones. **g.** enrichment of hypoallometric (left) and hyperallometric (right) ZRTscaling genes in cortical-type specific cell markers.^2^ Circle size denotes enrichment ratio, significant associations (p<0.05, hypergeometric test) are highlighted with black outline.

As expected, total cortical surface area increased exponentially between 21- and 38-weeks gestation (β_age_=0.054, p<0.001; **Figure 3c-d**).^93–95^ While cortical surface area was moderately greater in males compared to females (β_male_=0.011, p=0.002), this relationship did not change with age (p=0.946). At each vertex in the cortical surface mesh (n=30,248, excluding midline regions), we modelled areal expansion with respect to total surface area using log-log regression (see **Methods; Figure 3e**).^42^ Hyperallometric expansion, occurring at a rate faster than the cortical surface as a whole, was observed across the lateral neocortical surface encompassing the fronto-parietal operculum and (granular) insula, primary motor and sensory cortex as well as dorsal parietal and frontal cortices, confirming previous observations based on fetal MRI and scans of preterm-born infants (**Figure 3c, e**).^95–98^ In line with proposed models of cortical evolution and expansion,^5,9^ slower rates of growth were observed in medial allocortex (including entorhinal, paleocortex and parahippocampal cortex) and the cingulate cortex (**Figure 3e**). The inclusion of sex and sex:age interaction effects in the scaling model did not affect estimated vertex scaling coefficients (r = 0.996). We confirmed that estimates of cortical expansion from cross-sectional analysis aligned closely to longitudinal observations from a single fetus scanned three times during gestation (**Figure S11**).

We calculated the non-parametric correlation (Kendall’s *τ*) between regional estimates of ZRT gene expression in the cortical plate and subjacent tissue zones and average allometric scaling in each of cortical areas defined by the μBrain atlas (**Figure 3e**). In total, across both early and mid-gestation timepoints, expression of 433/2145 (20.1%) ZRT genes was spatially correlated with areal expansion during gestation in at least one tissue zone (ZRT_scaling_; n=542 significant associations, p_FDR_ < 0.01) (**Table S9, S10**). Associations with areal scaling were significantly more common in ZRT genes than in non-ZRT genes (ZRT: 20.1%, non-ZRT: 8.3%; odds ratio=2.78, p<0.0001) with most significant ZRT_scaling_ associations (414/542) localised to the CP (**Figure 3f; Table S10**). ZRT_scaling_ genes in the CP included known molecular correlates of areal identity (*EFNA5,*^99^ *GLI3*,^100^ *FGFR2*^101^) and axonal guidance (*SLIT1*, *ROBO3, SRGAP1*).^102^

Differential expression of ZRT_scaling_ genes largely captured differences between post-mitotic allocortex and neocortex, reflecting opposing allometric scaling across phylogenetic cortical types (**Figure 3e**). We found evidence at 15 PCW, but not at 21 PCW, that genes with higher expression in slower-expanding allocortex and peri-allocortex, were significantly enriched in early-born Cajal-Retzius neurons (e.g.: *CALB2*; overlap=17, enrichment=1.72, p_hypergeom_=0.021),^2^ cells that originate from the pallial-subpallial boundary and cortical hem and migrate tangentially across the developing neocortex in early gestation.^103,104^ ZRT_scaling_ genes involved in Notch signalling (*NOTCH2NLR*, *JAG1*)^105^ and others critical for hippocampal dendritic development (*LRIG1*)^106^ were also expressed highly in allocortical regions (**Table S10**). In contrast, ZRT_scaling_ genes expressed in the preferentially expanded neocortex were enriched in progenitor cells at 15 PCW (*FBXO32, HES6*; IPC enrichment = 1.51, p_hypergeom_ = 0.027), and general markers of deep layer neurons at both timepoints (*NEUROD6*, *SYT6;* 15 PCW: Neuron enrichment = 1.53, p_hypergeom_ = 0.004; 21 PCW: enrichment = 1.51, p_hypergeom_ = 0.007). While basic cell types are generally conserved across cortical areas,^107^ previous evidence has shown that regional identity is imprinted during cell differentiation, with areal signatures most apparent in post-mitotic cell types but pervasive even at early stages of development across major brain structures.^2,15,31^ In line with this, we found opposing enrichment of postmitotic neuronal markers specific to allocortex and neocortex in hypoallometric and hyperallometric ZRT_scaling_ genes, respectively (**Figure 3g**).

An expanded neocortex is a hallmark of the primate brain. A recent transcriptomic survey of the neocortex across primate species identified a set of genes differentially expressed in humans (hDEGS) and located near to genomic regions that are highly conserved across mammals but significantly altered along the human lineage, either through accelerated DNA substitution rates (human accelerated regions; HAR) or deletions (human conserved deletions; hCONDELS).^107–109^ We tested whether these genes were associated with human neocortical expansion *in vivo*. We found that ZRT_scaling_ genes were significantly enriched for hDEGs located near HARs (overlap = 37; enrichment=2.09, p_hypergeom_<0.0001) and hCONDELS (overap=17; enrichment=2.0, p_hypergeom_=0.008). Of these, 22 (56%) were expressed more highly in neocortical than allocortical regions, including several cell adhesion molecules (*DSCAM*, *PCDH7, PCDH9*, *LRFN2*), teneurins (*TENM3*) and ephrins (*EFNA5*), as well as genes with functional links to language acquisition (*FOXP2*) and neurodevelopmental disorders (*MEF2C*, *AFF2*, *ZEB2*) (**Table S9**).

## Prolonged neural migration precedes faster expansion across the neocortex

Focusing further on neocortical expansion, we removed allo- and transitory periallo-cortical structures (hippocampus, retrosplenial cortex, entorhinal cortex and paleocortex) and repeated our regional correlation analysis over all ZRT genes. Within the neocortex, a subset of 116 ZRT genes (including 113 ZRT_scaling_ genes) were significantly associated with differential rates of expansion across neocortical regions (ZRT_neo_; p_FDR_<0.01), with most associations localised to the intermediate zone (IZ; **Figure 4a; Table S10**). ZRT_neo_ genes were also enriched for hDEGS located near HARs (overlap= 10; enrichment=2.03, p_hypergeom_=0.028) including *PCDH7*, *PCDH9*, *TENM3* and *AFF2* but not hCONDELS (**Table S9**).

**Figure 4:**
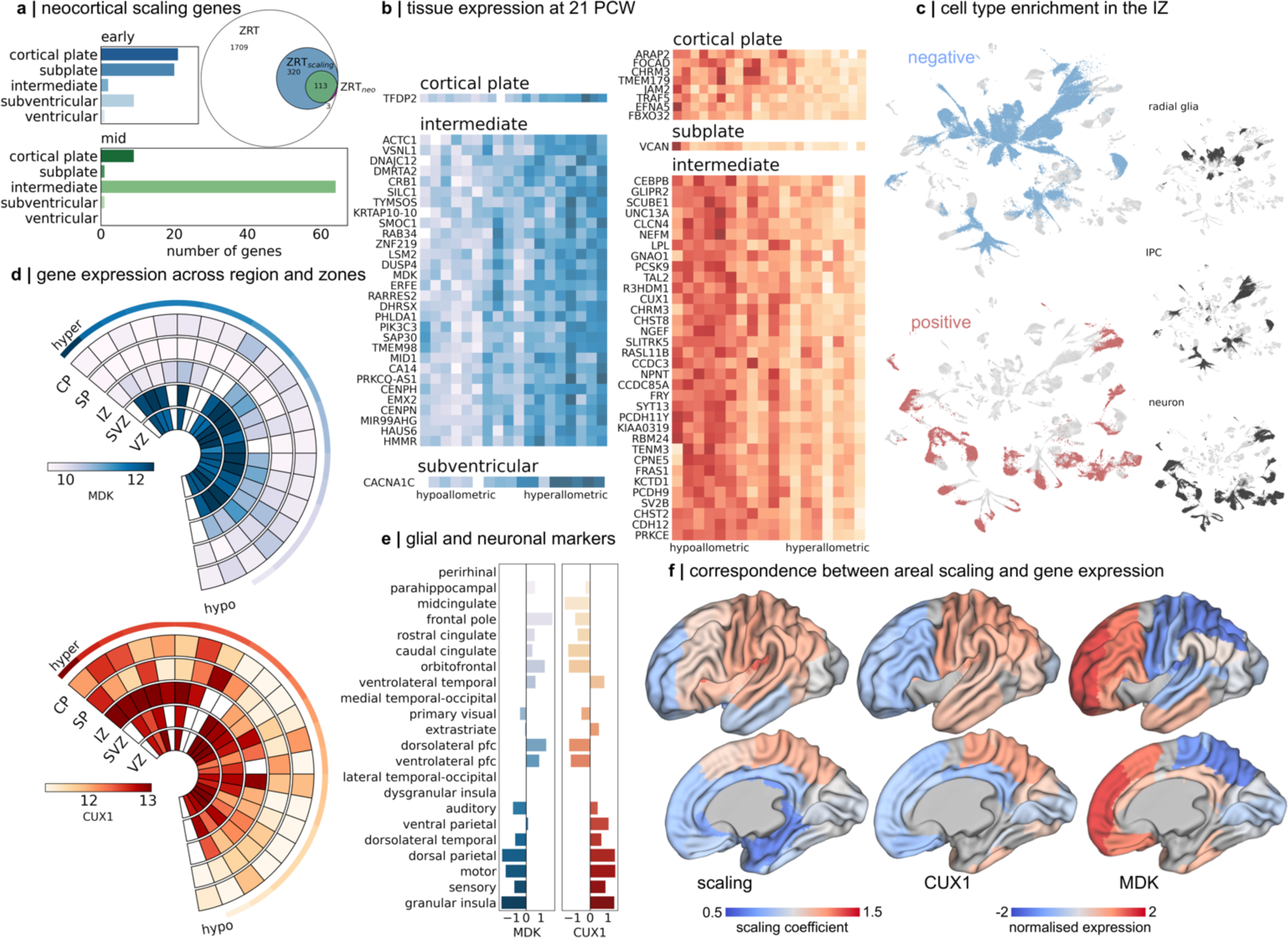
Preferential neocortical expansion is associated with differential timing of neurogenesis and gliogenesis. **a.** 133 ZRT genes were associated (pFDR<0.01) with areal scaling of the neocortex (after excluding paleo- and archi-cortex; ZRTneo). Most significant associations were localised to the IZ. **b.** normalised (Z-score) expression profiles for genes correlated with areal scaling in each tissue zone at 21 PCW. Associations at 15 PCW are shown in **Figure S12**. Negative associations (higher relative expression in hypoallometric regions) shown in blue, positive associations are in red. Lighter colours indicate higher relative expression. Most significant associations are in the IZ. **c.** Mid-gestation cell clusters^2^ significantly enriched (p<0.01) for genes associated with areal scaling in the IZ at 21 PCW. Territories of three cell types are shown. Negative and positive ZRTneo genes are enriched in progenitor cells and neurons, respectively **d.** wedge plots are shown for two ZRTneo genes expressed by specific cell types: *MDK* (glial) and *CUX1* (upper layer neurons). Rows indicate tissue zones and columns indicate cortical regions ordered according to allometric scaling from hyper to hypoallometric. Colour bar indicates normalised expression levels (a.u.). **e**. expression (Z-score) of *MDK* and *CUX1* in all regions sampled in the IZ, ordered from hypo (top) to hyperallometric (bottom) scaling. **f**. IZ expression of *CUX1* (middle) and *MDK* (right) projected onto corresponding μBrain surface atlas labels and displayed on the 36w dHCP template surface. Regions where expression for a given gene was not available are shown in grey. For comparison, average allometric scaling in each region is displayed (left).

We observed contrasting cell type enrichments of ZRT_neo_ genes at 15 and 21 PCW. Consistent with role of prolonged radial glial proliferation in proposed models of cortical expansion,^11,51,54^ highly expressed ZRT_neo_ genes in areas with a higher rate of expansion over gestation were enriched in radial glia and intermediate progenitors at 15 PCW (p_hypergeom_=0.045, 0.040 respectively; **Figure S12; Table S11**) with significant associations localised to the cortical plate, subplate and subventricular zone (**Figure 4a**). Early hyperallometric ZRT_neo_ genes are upregulated in both outer (*CDC42EP4*, *HS6ST1*) and ventricular (*FBXO32*) radial glial subpopulations^53^ (**Figure S12**). In contrast, ZRT_neo_ genes expressed in neocortical areas with slower relative growth were localised to the cortical plate and subplate but not specifically enriched for any major cell types (all p_hypergeom_>0.05; **Table S11**). However, hypoallometric ZRT_neo_ genes were expressed by neurons (*NFE2L*) and involved in dendritic (*ABGRB3*^110^) and synaptic (*NPTX2*^111^) plasticity, indicative of a population of maturing, not proliferative, cells with neuronal lineage in these regions.

At 21 PCW, after the peak period of neurogenesis, we observed the opposite pattern of cell type enrichments. ZRT_neo_ genes expressed in the IZ subjacent to preferentially expanded cortical areas were enriched in neuronal populations (enrichment = 2.19, p_hypergeom_= 0.00011) (**Figure 4a-c**) whereas, hypoallometric ZRT_neo_ genes were enriched in proliferative glial cell types (IPC: enrichment=2.96, p_hypergeom_ <0.0001; RG: enrichment=2.36, p_hypergeom_<0.0001; **Figure 4b,c; Table S11**). The presence of post-mitotic neuronal markers in the IZ at 21 PCW suggested that neuronal migration was ongoing in cortical areas with the fastest rate of expansion in the third trimester of gestation. This is consistent with a conserved mechanism of mammalian cortical expansion whereby longer neurogenic periods lead to an expanded neocortex.^11,112–115^ In this context, on both phylogenetic and ontogenetic scales, later developing cortical regions would exhibit faster rates of expansion.^51,115,116^ A prominent hypothesis of neocortical expansion has suggested that, in primates, this process is realised through the continued production of upper layer neurons from outer radial glia (oRG) populations situated in the outer SVZ, a cell population greatly expanded in the primate brain.^51,54^

To examine this proposed mechanism in humans, we focused on *CUX1*, a marker of layer III/IV neurons that regulates dendritic morphology^117^ and is expressed highly in preferentially-expanded cortical regions (**Figure 4d-f**). *CUX1* is located downstream of HAR426 and pathogenic mutations in *CUX1* are associated with ASD, intellectual disability and epilepsy.^118,119^ We find that, in the IZ at 21 PCW, *CUX1* exhibits expression that varies along a hypo-to-hyperallometric gradient (**Figure 4d,e**; *τ*=0.52, p_FDR_=0.002). To validate these observations, we examine ISH staining of a second upper layer marker, *SATB2*, in five regions with differential allometric scaling, finding examples of upper layer SATB2^+^ neurons within the IZ of regions with a faster rate of expansion in mid- to late-gestation (**Figure S13**). The prolonged migration of these cell populations in expanding neocortical regions is a potential consequence of differential neurogenic timing across the neocortical sheet that, at least in part, supports the accelerated expansion of hyperallometric cortical regions during gestation.

Several mechanisms exist to regulate gene transcription during early brain development.^120,121^ To identify potential regulators of ZRT gene expression in the developing fetal cortex, we used a recent chromatin accessibility atlas^122^ to examine the position of open chromatin regions (OCR) in the mid-gestation brain relative to ZRT genes. We found that ZRT genes were more likely than non-ZRT genes to be located near to predicted regulatory elements (pREs), a subset of OCRs that are likely to function as neurodevelopmental enhancers in mid-gestation^122^ (OR: 1.38 p < 0.0001; **Figure S14**; **Table S12**). Moreover, ZRT_scaling_ and ZRT_neo_ genes were significantly enriched for genes located near to pREs (enrichment = 1.25, 1.31 p_hypergeom_<0.0001, <0.005 respectively; **Table S12**). Focusing on laminar specificity of ZRT gene expression, we found that over 25% of ZRT_neo_ genes were located immediately up- or downstream of OCRs specific to the upper layers of the cortical plate, compared to 9% located near to deep layer OCRs (**Figure S14B**). Transcription factor motifs contained within OCRs specific to upper cortical layers and proximal to ZRT_neo_ (n=20) included bHLH, LIM and POU homeobox and HMG-box motif families (**Figure S14C**) that bind to transcription factors which regulate superficial neuronal identify (e.g.: *E2A*, *BRN1*, *LHX2*).^123–125^ Thus, the differential accessibility of specific regulatory elements can resolve the laminar identity of maturing upper-layer excitatory neurons migrating through the IZ at 21 PCW.

Based on this evidence, we reasoned that neuronal migration, and thus neural proliferation, would be complete or near complete at 21 PCW in neocortical areas with slower expansion rates in the third trimester. In this case, expression of proliferative cell markers (**Figure 4**) would reflect gliogenesis rather than neurogenesis. To test, we compared ZRT_neo_ genes associated with cortical scaling at 21 PCW in the IZ to region-specific cell type signatures in the mid-fetal brain.^2^ Reflecting the proximity to medial allocortex and periallocortical regions, we identified several midline identity genes (*MID1*, *DMRT5*, *EMX2*) with high expression in hypoallometric cortex as well as markers of cell proliferation (*HMMR*, *HAUS6*, *CENPN*, *CENPH*, *PIK3C3*) (**Figure 4b; Table S10**). In support of our hypothesis,, we found that hypoallometric ZRT_neo_ genes were specifically enriched in (peri)allocortical glial cell populations (*MDK*, *SAP30*, *TMEM98;* astroglia, p_hypergeom_=0.01; OPC, p_hypergeom_=0.07).

TMEM98 is a MYRF-interacting protein specifically expressed in newly-differentiated oligodendrocytes in the developing central nervous system^126^ whereas MDK is a growth factor expressed in pre-OPCs that can induce differentiation in oligodendrocyte-lineage OL1^+^ cells *in vitro*.^127–129^ *SAP30* forms a co-repressor complex with *HDAC1* and *HDAC2*, class I histone deacetylases that regulate gene transcription and are essential for oligodendrocyte maturation.^130–133^ Similar negative correlations with cortical expansion were recorded in OPC cell population markers S100B (**Table S10**; *τ*=-0.33, p_FDR_=0.070), NKX2-2 (*τ*=-0.41, p_FDR_=0.019) and the glial progenitor marker *EGFR* (**Figure S15**), which has been validated previously as a critical gene related to brain size.^134^ In an independent dataset,^33^ we observed similar trends in *OLIG1* expression in mid-gestation across cortical regions with differential developmental expansion (**Figure S16**). Overall, these data suggest that the developmental timing of the neuro- to gliogenic switch varies across the neocortical sheet, with the length of the neurogenic period supporting differential rates of neocortical expansion during the third trimester of gestation.

## Neocortical scaling genes are critical for typical neurodevelopment

Given their likely importance in shaping early normative neurodevelopment, we hypothesized that the ZRT_neo_ genes would be susceptible to severely disruptive mutations (i.e., loss-of-function variants). We found significant enrichment of hyperallometric (median loss of function observed/expected upper bound fraction (LOEUF) score = 0.26, permutation p = 0.0003 using random gene sets of similar size: p_permutation_) but not hypoallometric (median LOEUF score = 0.40, p_permutation_ = 1) ZRT_neo_ genes, suggesting a disproportionate level of vulnerability to loss-of-function variation in genes whose expression is greater in areas that expand fastest in the third trimester. Within these, we identified a set of constrained genes expressed highly in the subventricular zone at 15 PCW in hyperallometric regions. These genes are involved in extracellular matrix formation and interaction (*EFEMP2*, LOEUF=0.56, *PTPRM*, LOUEF=0.33), and epithelial-to-mesenchymal transition (*FBXO32*,^135^ LOEUF=0.64), pathways crucial to outer radial glia specification and differentiation in germinal zones of the developing brain.^53^ Follow-up analyses using genome-wide metrics for dosage sensitivity^136^ confirmed the enrichment of hyperallometric ZRT_neo_ genes as haploinsufficient (62% of genes, p_permutation_ < 0.0001 using random gene sets of similar size) and not triplosensitive (19%, p_permutation_ = 0.9418) – a highly pathogenic mechanism for loss-of-function mutations.

To assess the clinical relevance of these distinct ZRT_neo_ gene sets (i.e., hypoallometric and hyperallometric), we performed enrichment analyses using MAGMA^137^ across an array of previously published genome-wide association studies (GWAS). We found that ZRT_neo_ gene sets were not enriched for birth outcomes (gestational duration) or cognition (educational attainment), but hypoallometric ZRT_neo_ genes were enriched for externalizing behavior (β=0.17, p= 0.007) and hyperallometric ZRT_neo_ genes were enriched for schizophrenia (SCZ; β=0.17, p=0.004). Further analysis using postmortem gene expression data from patients with neurodevelopmental disorders revealed significant enrichment of ZRT_neo_ gene sets within multiple co-expression modules.^138^ Both hypoallometric and hyperallometric ZRT_neo_ genes were enriched in cross-disorder module *CD1* (both p_permutation_ < 0.05) – downregulated in autism spectrum disorder (ASD), SCZ, and bipolar disorder, and containing neuron-enriched genes and genes with ASD- and SCZ-associated nonsynonymous de novo variants from whole-exome sequencing; and hyperallometric ZRT_neo_ genes were enriched in module *CD13* (p_permutation_ < 0.05) – also downregulated in ASD, SCZ, and bipolar disorder, and containing neuron-enriched genes.

## Discussion

Despite the altriciality of the human brain at birth, areal expansion of the cortex during the second and third trimester of gestation is critical for later neurodevelopmental function. Cortical surface area increases exponentially during the third trimester of gestation, permitted by rapid cortical folding over the same period. Powered by a new 3D atlas of the developing brain, our results provide a multiscale understanding of fetal cortical expansion in the second half of pregnancy. We find that differential expansion of cortical areas in gestation respects anatomical and evolutionary boundaries between cortical types^5,9^ and is supported by an extended period of neural migration through mid-gestation.^51,52,54^

Neurogenesis exhibits a conserved order but nonlinear scaling across species.^139^ Longer neurogenic periods in larger-brained species, supported by a larger pool of progenitors in proliferative zones, result in the preferential expansion of later developing structures.^11,112–114,139^ In mammals, differences in the timing and rate of neuron production vary across cortical areas with evidence to suggest that progressive termination of cortical neurogenesis occurs along a rostral-caudal axis.^140–142^ In this case, earlier termination of neuronal production in the anterior cortex could create a potential affordance for increased neuronal size and arborisation, leading to increased areal expansion during development.^25,139,143^ However, further evaluation of areal differences in neurogenic timing in the primate cortex presents a more complex picture, with neurogenesis terminating first in limbic and allocortical structures but continuing in the prefrontal cortex beyond mid-gestation.^139,144,145^ Coupled with the nonlinear progression of human gyrification over gestation,^146^ this suggests that areal differences in cortical scaling are likely founded upon an alternative schema.^49,51^

Alternative hypotheses have been put forward on the role of oRG proliferation, and the prolonged production of neurons or glia, in cortical expansion.^49–52^ Our findings demonstrate that prior to gyrification, but after the peak period of neurogenesis, supragranular neurons continue to migrate to neocortical areas with the fastest rate of expansion in the third trimester. In the primate brain, oRGs produce large numbers of upper layer neurons, provide a scaffold for neural migration and, upon completion of neurogenesis, act as a source of glial cells in mid- to late-gestation.^49,52,54,147^ Thus, regulation of neuro-to-gliogenic timing in the oRG subpopulation may represent a plausible candidate for differential rates of neocortical expansion^49,147^ Though present in other mammals, the oSVZ is expanded in primate species^51,53,54^ and proliferation in the oSVZ, marked by mitotic activity, is highest in regions that expand most in later development.^148^ While our data suggest rapid areal expansion is preceded by an extended neurogenic period, we lack the data to confirm a similarly extended period of gliogenesis. In humans, neurogenesis precedes cortical folding with the subsequent gliogenic period more closely aligned to the timing of cortical expansion.^49,50^ An extended neurogenic period coupled with a longer migration time due to the expanding volume of the brain may necessitate an extended gliogenic period to populate the expanding neuropil.^50^ Evaluating the temporal and spatial regulation of glial fate transition and proliferation in the oSVZ during the second half of gestation represents a critical next step in understanding this process.

In gyrencephalic species, the buckling and folding of the cerebral cortex allows for increased surface area of the cortical grey matter. Greater tangential expansion of superficial cortical layers relative to subcortical tissue represents a core feature of biomechanical models of cortical growth and folding.^149–152^ However, uniform rates of tangential expansion can not fully account for the consistency in location of cortical folds across individuals, with additional genetic contributions to gyral patterning clearly demonstrated in twin studies.^153,154^ In contrast, genetically determined areal differences in expansion rate may give rise to the consistent patterns of folding observed across the neocortical sheet.^148,155^ Recently, large-scale neuroimaging studies have identified patterns of altered cortical morphometry that are shared across common neuropsychiatric conditions and human genetics studies have begun to converge on putative mechanisms underlying cortical abnormalities in developmental genetic disorders.^156–160^ Here, we identify significant enrichment of pathogenic loss-of-function variants in genes that are expressed in mid-gestation, linked to specification of outer radial glia and associated with differential rates of cortical expansion. Taken together, these findings suggest that there are temporal windows of susceptibility in the early stages of brain development where areal differences in the timing of fundamental neurogenic processes could underlie observable cortical abnormalities and postnatal functional pathologies in neurogenetic disorders.^47^

Spatially-embedded gene expression atlases of the adult human^35,68^ and mouse^70,161^ have proven exceptionally powerful in recent years, bridging resolution gaps to common neuroimaging modalities^162,163^ and providing insight into the molecular correlates of structural^26,42,164^ and functional neuroanatomy,^36,165,166^ brain development,^167–169^ disease and disorder.^47,170,171^ In such studies, comparisons with *in vivo* neuroanatomy can only be fully realised through three-dimensional localisation of tissue samples within a common coordinate space.^62,68,70^ To date, a limitation of this approach has been either the sampling of a narrow age range outside of key developmental periods^35,69^ or, in developmental datasets, a lack of 3D spatial information^3^ and relatively coarse anatomical sampling.^34^ To fill this gap, we provide a new resource, μBrain, built upon existing open-source data, to allow researchers to map developmental neuroanatomy of the human fetal brain onto early histogenic processes using contemporaneous *post mortem* data. The reconstructed 3D μBrain atlas brings detailed tissue microarray and *in situ* hybridisation data into alignment with a developmental anatomical atlas of the fetal brain.^89^ The μBrain atlas will enable future studies to examine tissue or region-specific expression signatures in relation to aspects of structural or functional brain development *in utero* or identify spatial or temporal windows of vulnerability for genetic or neurodevelopmental disorders.

Developmental MRI studies provide unique insight into early human brain development. Due to large differences in size, shape and tissue contrast, specialised tools are required for the analysis of infant and neonatal MRI. Similarly, we cannot rely on common cortical atlases that are based on adult neuroanatomy.^87,172^ Here, we used annotations derived from cytoarchitecture of the mid-fetal brain to generate a new cortical atlas to facilitate further research in early brain development. A key area for future research in this field is the development and validation of improved methods to align early MRI to common template spaces. The geometry of the fetal cortex is smooth, making alignment of cortical morphometry an ill-posed problem. Newer, anatomically-constrained registration techniques and larger longitudinal cohorts with multiple scans during mid- to late-trimester will enable more precise estimates of cortical expansion in the future.^90,95^

With increasingly granular surveys of the developing brain at a single-cell level^2,107^ the advent of spatial transcriptomic technologies,^173^ and a series of large-scale and open-access perinatal neuroimaging studies,^84,174,175^ we anticipate μBrain will provide a foundation for developmental and comparative neuroscience to integrate and transfer knowledge of early brain development across domains, model systems and resolution scales.

## Acknowledgements

This research was supported by the National Health and Medical Research Council (NHMRC) [1194497 to G.B.], the Murdoch Children’s Research Institute, the Royal Children’s Hospital, Department of Paediatrics, The University of Melbourne and the Victorian Government’s Operational Infrastructure Support Program. The project was generously supported by RCH1000, a unique arm of The Royal Children’s Hospital Foundation devoted to raising funds for research at The Royal Children’s Hospital. L. Z. J. W was supported by the Commonwealth Scholarship Commission, United Kingdom. V. Ka. was supported by an MRC (UK) award MR/V036874/1 and The Developing Human Connectome Project. E.C.R. was supported by a Wellcome Collaborative Award (215573/Z/19/Z). AA-B and JS were supported by R01MH132934.

Neuroimaging data were provided by the Developing Human Connectome Project, KCL-Imperial-Oxford Consortium funded by the European Research Council under the European Union Seventh Framework Programme (FP/2007-2013)/ ERC Grant Agreement no. [319456]. We are grateful to the families who generously supported this trial.

We are grateful to the Allen Institute and associated Investigators for the provision of the Atlas of the Developing Human Brain.

## Author contributions

G.B., S.O. V.K., L.Z.J.W., V.K., A.P., J.V.H., J.H., E.C.R. & J.S. performed data acquisition and data processing. G.B., S.O. & J.S performed data analysis. G.B, J.S., V.K., L.Z.J.W & E.C.R. contributed to methodology. G.B., M.L.S., A.A-B., J.V.H., A.D.E., E.C.R. & J.S. provided resources and supervision. E.C.R. provided software. Project conceptualisation: G.B & J.S. Writing drafts, revisions and editing: all authors.

## Material and Methods

### Public data sources

Source data underlying the μBrain atlas were made available as part of the BrainSpan Developing Brain Atlas [https://atlas.brain-map.org/atlas?atlas=3] with detailed tissue processing protocols available from Ding et al.^3^ In brief, a single prenatal brain specimen (21 PCW; female) was bisected and the right hemisphere used for serial sectioning. The brain specimen was cut into four coronal slabs and frozen in isopentane. Serial coronal sectioning at 20μm thickness was performed slab-by-slab with sequential sections submitted to Nissl, AChE or ISH staining with 43 gene probes and stained sections digitally scanned at 1μm/pixel resolution. In total, 81 out of 174 Nissl-stained sections with varying sampling densities (∼0.5mm to 1.2mm between sections) were selected for annotation.^3^ Expert anatomical annotations were conducted manually on each section. Nissl- and ISH-stained sections with corresponding anatomical labels were made available for download. Anatomical annotations were also used to guide laser microdissections for DNA microarray analysis across the developing cerebral tissue in the left hemisphere of 4 separate mid-gestation specimens (see **Microarray Data** below). The section numbers and approximate coronal positions of sections used in the construction of the 3D μBrain atlas are listed in **Table S1**.

### Image processing

We downloaded each high-resolution Nissl-stained section (n=81; downsampled to 2μm/pixel) as RGB images in JPG format with corresponding anatomical labels as SVG files.

After converting SVG to RGB PNG format, we manually combined anatomical labels according to the hierarchical ontology of the reference atlas^3^ to create two compact annotations, one for image repair comprising 20 tissue structure labels (*brain-labels*) and one for statistical analysis containing only cortical labels (*cortex-labels*, n=30, including one generic ‘brain tissue’ label for non-cortical structures; see **Table S2**). Due to the small size and degree of missing data precluding reconstruction, marginal zone and subpial granular zones were not considered in this analysis. Nissl-stained sections and corresponding label images were then downsampled to 20μm/pixel resolution.

### Histological reconstruction

*Pix2pix* is a conditional generative adversarial network (GAN) trained to perform image-to-image translation between pairs of image examples.^59^ We used the *pix2pix* architecture (**Figure 1b**) to synthesise Nissl-stained images from label annotations in order to replace artefacts within tissue sections (**Figure 1c,d**). Following conventional GAN structure, the model combines a generator network, *G*, with a classifier (or discriminator, *D*) whose objective is to determine if images are real or fake (**Figure 1b**). GAN training is performed adversarially with the generator network competing to generate more and more realistic synthetic image from label annotations, and the discriminator working to discriminate between real and fake examples. An *L*_1_ regularisation term is added to enforce that generated images are as close as possible to the ground truth. Full model architecture and training details are included in the **Supplemental Methods**.

### Image repair

To perform repair of whole sections, we split each label image into patches of 256 × 256 pixels with an 8 pixel overlap and passed them through the trained generator. The resulting, synthetic Nissl contrast patches were stitched together into a full section matching the dimensions of the original image (**Figure 1d**). Patch prediction and image reconstruction was performed using MightyMosaic [https://pypi.org/project/MightyMosaic/].

To detect regions of the original Nissl-stained section that needed repair, we designed an automated outlier detection method based on the Median Absolute Deviation (MAD) of pixel hue and saturation. The original Nissl-stained sections and corresponding GAN-generated predictions were transformed to HSV format and blurred with a box filter (width = height = 5 pixels). We identified outliers with median absolute differences in hue and saturation between pixels in the ground truth image and its synthetic equivalent in hue and saturation greater than threshold, *θ*, set to 2.5, whereby lowering *θ* would increase the number of pixels marked as outliers.

For each section, a binary mask was created containing all pixels identified as outliers in both hue and saturation. A final opening operation was applied to the outlier mask using an elliptical filter (iterations = 3, width = 3 pixels) to remove speckles in the mask. Identified outlier pixels were then replaced with the corresponding, intensity-matched pixels from the synthetic image using Poisson image editing to effect image repair (Figure 1d).^61^ Outlier detection and repair was performed in Python using OpenCV (4.5.2) [https://opencv.org/].

### μBrain volume construction

Following automated repair of major tissue artefacts present in the histological data, we aimed to develop a 3-dimensional reconstruction of the fetal brain to facilitate comparison with *in vivo* MR imaging data. Image alignment and reconstruction steps are summarised below. Full details are included in **Supplemental Methods**.

#### Slice-to-slice alignment

Using the middle section as a reference, repaired Nissl-stained sections were aligned using a graph-based, slice-to-slice registration.^176,177^ Pairwise rigid transforms were estimated between each section and its neighbouring sections in the direction of the reference. Dijkstra’s shortest-path algorithm was then used to calculate the set of transforms with lowest cost to align a given section to the reference.^176,177^ The selected transforms were composed and applied to both the image and its corresponding labels to bring all sections into approximate alignment (**Figure 1e**; **Figure S17a**).

#### Affine registration to a fetal brain shape reference

Reconstructing 3D volumes from the consecutive alignment of 2D sections commonly produces an artefact termed ‘z-shift’ caused by the propagation of registration errors between adjacent slices and resulting in a distorted three-dimensional structure in the final volume.^178^ To overcome this effect, it is common to use a shape prior to guide registration and preserve 3D shape.^62,178,179^ In lieu of a ground-truth volume for the sectioned data, we employed a population-based average anatomical image: specifically the 22-week timepoint of the Gholipour et al. spatio-temporal fetal MRI atlas (**Figure S3**).^63^

After matching MRI-based tissue labels to the μBrain tissue labels, we upsampled the MRI template to 50μm isotropic resolution and converted the MRI labels into an image Nissl-like contrast using the trained GAN model (**Figure S3c-d**). Nissl-contrast images were re-stacked into a 3D volume to act as an anatomical prior for registration.

We performed an iterative affine registration procedure between the MRI-based shape prior and the 3D stack of histological sections.^176^ This process was repeated for a total 5 iterations, producing a final 3D volume with aligned coronal slices and a global shape approximately matched to the *in utero* fetal brain (**Figure 1e; Figure S17a**).

#### Final template construction

To create the final 3D volume, we employed a data augmentation technique, generating n=50 unique representations of the affinely-aligned data by applying nonlinear distortions along all three image axes. For each volume, we performed a weighted nonlinear registration between neighbouring sections to account for residual misalignments. Finally, to create a smooth 3D reconstructed volume, we co-registered all 50 augmented and aligned volumes into a single probabilistic anatomical template with voxel resolution 150 × 150 × 150μm using an iterative, whole-brain nonlinear registration (**Figure 1e**; **Figure S17a**; **Supplemental Methods**). All image registration was performed in Python 3.7 using *antspyx* (0.2.7).^14^

#### Cortical reconstruction

To reconstruct the fetal cortical surface, we adapted existing protocols for *ex vivo* [https://freesurfer.net/fswiki/ExVivo] and non-human primate [https://prime-re.github.io/] surface reconstruction with Freesurfer.^15^ We used the μBrain tissue labels to generate a ‘white matter’ mask (all subcortical structures and tissue zones, excluding the cortical plate). We used this mask to generate inner and outer surfaces for the μBrain volume (**Figure 1f**). Surfaces were smoothed and inspected for topological errors. All processing was performed with Freesurfer (7.3.2).

### In situ hybridisation

In addition to serial Nissl staining, interleaved coronal sections were used for *in situ hybridisation* (ISH) of a series of neurodevelopmental marker genes (**Table S3**).^3^ High-throughput ISH staining was performed for each gene, with stained sections digitised at 1*µ*m resolution. Quantification of the intensity of expression detection was performed using an automated procedure that pseudo-colour coded levels of expression for visualisation, with low-to-high expression represented as blue-to-red.^161^

Compared to Nissl-stained sections (n=79 after quality control), fewer ISH stained sections were available for each gene (mean n = 41 after quality control), precluding a full 3D reconstruction of each. We downloaded each set of ISH-stained sections and removed any with large artefacts (tearing, folding, missing tissue). From each false-colour expression map, we extracted the red channel to focus only on higher expressing cells. Each section was registered to the nearest, repaired Nissl-stained section using affine registration. Registrations were visually inspected and any failures removed. Aligned sections were then stacked together, with blank slices in place of missing sections and reconstructed into a 3D volume using the previously calculated slice-to-volume alignments for each section (see ‘**μBrain volume construction**’).

### Microarray data

We downloaded prenatal LMD microarray data from the BrainSpan database [https://www.brainspan.org/]. For details on tissue processing and dissection see Miller et al.^4^ and the technical white paper available at: [https://help.brain-map.org/download/attachments/3506181/Prenatal_LMD_Microarray.pdf]. In total, normalised microarray data from 58,692 probes in 1206 tissue samples were available to download, obtained from the left hemisphere of four post-mortem fetal brain specimens (age 15-21 PCW, 3 female).^4^ Each probe was assigned a ‘present’ or ‘absent’ annotation based on strength of average probe expression over corresponding background signal. Through comparison with the BrainSpan reference atlas, we matched each tissue sample’s anatomical label to i) corresponding cortical labels included in the μBrain atlas and ii) one of five tissue zones (cortical plate, subplate, intermediate zone, subventricular zone, ventricular zone) (**Table S4; Figure S4**). Samples that could not be matched to labeled regions in the cortical plate or corresponding subjacent tissue zones were removed, including samples from subcortical nuclei, midbrain structures and brainstem.

#### Microarray processing

We updated gene assignments for the Allen microarray probes using Re-Annotator^180^ and removed any probes assigned to more than one gene, resulting in a probe set (n=46,156) mapped to 20,262 unique genes. Low signal probes designated ‘absent’ were removed (34.67% of probes), as were tissue samples from the marginal zone, subpial granular zone and subcortical and midbrain structures (54.46% of samples). Where multiple probes mapped to a single gene, the probe with the highest differential stability (DS),^181^ the average pairwise correlation between tissue sample expression over all specimens, was assigned. Probes with DS<0.2 were removed.

Where more than one sample was available for a given region or zone, e.g.: samples from the outer and inner cortical plate in the same region, gene expression was averaged across samples. Finally, any probes with missing data in more than 10% of tissue samples were removed (n=1253). This resulted in expression data from 8771 genes across 27 regions and 5 tissue zones for analysis (**Figure S4**).

### Fetal MRI

To measure cortical expansion *in utero* during the third trimester, we analysed high-resolution MRI from a large cohort of fetuses.

#### MRI acquisition

Fetal MRI datasets (n=240 scans from 229 fetuses aged between 21^+1^ and 38^+2^ gestational weeks^+days^) were acquired as part of the Developing Human Connectome Project (dHCP) using a Philips Achieva 3T system, with a 32-channel cardiac coil in maternal supine position. Structural T1-weighted (T1w), T2w, functional MRI and diffusion MRI data were acquired for a total scan time of approximately 45 minutes.^85^ T2-weighted SSTSE volumes were acquired with TE=250ms, acquisition resolution 1.1 x 1.1mm, slice thickness 2.2mm, -1.1mm gap and 6 stacks. All 3D brain images were reconstructed using a fully automated slice-to-volume reconstruction (SVR) pipeline^86^ to 0.5mm resolution and reoriented to the standard radiological space.

The study was approved by the UK Health Research Authority (Research Ethics Committee reference 452 number: 14/LO/1169) and written parental consent was obtained in every case for imaging and open data release of the anonymized data. All data was acquired at St Thomas Hospital, London, United Kingdom.

After image processing and quality control, the final dataset comprised n=195 fetal MRI datasets acquired from n=190 fetuses aged 21^+1^ to 38^+2^ gestational weeks (88 female). Repeated scans were acquired from four fetuses.

#### MRI processing

While neonatal protocols for automated MRI tissue segmentation exist,^87,182^ due to the differences in size, tissue contrast and signal-to-noise ratio, segmentations derived from fetal MRI often require extensive manual editing to ensure accuracy.^183^

Here, we used an optimised neonatal tissue segmentation pipeline (Draw-EM)^87^ with tissue priors adapted to a fetal MRI template to create a ‘first-pass’ tissue segmentation for each fetal MRI volume. Tissue segmentations were then visually checked and extensive manual corrections performed where needed to correct gross segmentation errors and ensure accuracy of tissue boundaries (CSF/cortex/white matter). Manually-corrected tissue segmentations were then used to generate anatomically and topologically correct inner and outer cortical surfaces using *Deformable*.^88^ Note that all intensity-based correction terms were turned off during surface reconstruction and each surface was generated using just the corrected tissue segmentations. At each stage, images and derived outputs were visually inspected for accuracy.

#### Alignment to fetal template

We aligned individual cortical surfaces to the dHCP fetal atlas, a spatiotemporal surface atlas, spanning 21-36 weeks of gestation with weekly timepoints.^89,91^ Using MSM with higher-order clique reduction, we calculated non-linear transforms of individual surfaces to their closest fetal timepoint based on spherical registration of sulcal depth features.^90,92^ The MSM transform was used to resample individual surface topology (pial, midthickness, and white) onto the template surface vertices, ensuring that all surfaces across individuals had the same vertex correspondence. Resampled surfaces were manually checked to ensure the quality of the registration.

#### Alignment to μBrain

We aligned the μBrain cortical surface to the earliest timepoint of the dHCP fetal template surface using a two-step nonlinear surface registration guided by a set of anatomical priors (**Figure S17b,c**). We used MSM to perform an initial nonlinear spherical registration between μBrain and dHCP surfaces based on alignment of sulcal depth. After this, we created a set of coarse cortical labels on the dHCP surface matched to corresponding μBrain labels by combining a) dHCP cortical atlas labels,^87^ b) manual labels guided by sulcal anatomy on the 36 week fetal surface and c) combining μBrain labels in the same lobes (e.g.: ventrolateral frontal, dorsolateral frontal, orbitofrontal) were into single anatomical labels. The full list of 11 matched cortical regions included: auditory cortex; cingulate cortex; frontal cortex; insular cortex; primary motor; primary sensory; occipital cortex; parahippocampal cortex; parietal cortex; superior temporal cortex; ventrolateral temporal cortex. A secondary multivariate spherical registration between μBrain and fetal surfaces was initialised using the previously calculated sulcal alignment and driven by alignment of cortical ROIs across surfaces.^90^ This approach leverages anatomical labels (defined based on cytoarchitecture, or using older fetal anatomy in μBrain and dHCP atlases, respectively), to inform cortical alignment in the absence of geometric features. A similar approach has proven successful accommodating large deformations across primate species.^184^

μBrain labels were propagated to each timepoint of the dHCP fetal atlas (**Figure 3b**) and onto the surface topology of each fetal scan. Cortical labelling was visually quality checked for alignment.

### Statistical analysis

#### Allometric scaling of cortical surface area

Each subject’s outer cortical topology was resampled onto the dHCP template surface (32,492 vertices) and vertex-wise estimates of cortical surface area were corrected for folding bias by regressing out cortical curvature^185,186^ and smoothed with a Gaussian kernel (FWHM = 10mm). Total cortical surface area was calculated as the sum of all vertices in the cortical mesh, excluding the medial wall. At each vertex, *β*, we modeled scaling relationships with brain size by estimating the log-log regression coefficient for total surface area as a predictor of vertex area, *a_v_*:^42^

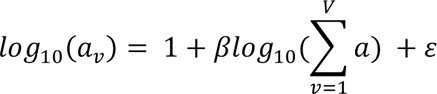

Such that the scaling coefficient, *β*, can be directly interpreted relative to 1 (representing linear scaling between vertex area and total area) with *β* > 1 and *β* < 1 representing hyper- and hypoallometric scaling of vertices with respect to total area, respectively. Models were fit using Ordinary Least Squares (OLS) regression. We tested alternative models including sex and age:sex interactions. Analyses were repeated after removing repeated scans to satisfy i.i.d. assumptions of OLS regression (n=190; **Figure S18**).

Prior to analysis, vertexwise outliers were identified and removed (**Figure S19**). To account for age-related increases in area, outliers were identified using a sliding window over age (outliers >2.5 S.D. from the mean within a given window, maximum window size=25 scans, sorted by age). Data from five scans were removed prior to analysis due to the presence of outliers in more than 5% of vertices.

Vertexwise maps of areal scaling (*β* coefficients) were parcellated using the μBrain cortical labels, calculating average scaling within each parcel for further analysis.

#### Modelling changes in gene expression over zone (Z), region (R) and time (T)

For each gene (n=8771), we modelled the main effects of cortical tissue zone, region and timepoint on expression using a general linear model. Significant effects (p<0.01) were identified after False Discovery Rate correction for multiple comparisons over genes. Statistical analysis was performed in *statsmodels* (0.13.5)

#### Enrichment analyses

For all enrichment analyses, we calculated the enrichment ratio as the ratio of the proportion of genes-of-interest within each geneset/marker list to the proportion of background genes within each geneset. Unless otherwise stated, the background set was defined as the full list of genes included in the study (n=8771). Significance was determined using the hypergeometric statistic:

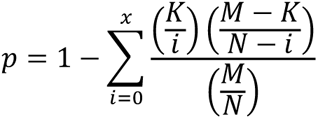

Where *p* is the probability of finding *x* or more genes from a specific geneset *K* in a set of randomly selected genes, *N* drawn from a background set, *M*. Where stated, False Discovery Rate (FDR) correction was applied to multiple comparisons.

### Code and data availability

The μBrain digital template with corresponding cortical surfaces and atlas labels is available from https://garedaba.github.io/micro-brain alongside code supporting data processing and analysis for this manuscript.

All dHCP data, fetal brain reconstructions, brain region segmentations and cortical surfaces are available for download from the NDA https://nda.nih.gov/edit_collection.html?id=3955

Source histological and microarray data are available from the Allen Brain Institute https://www.brainspan.org/

