## Supplemental Information for "Molecular signatures of cortical expansion in the human fetal brain"

### SUPPLEMENTAL MATERIALS

**Supplemental methods**

**GAN model architecture and training**
Model architecture
Training data
Model training
Model evaluation

**Template construction**
Slice-to-slice alignment
Affine registration to a fetal brain shape reference
Final template construction
Cortical reconstruction

**Supplemental figures**

Figure S1: Lightbox of reference atlas sections
Figure S2: Model performance over hyperparameters
Figure S3: Shape reference for affine registration
Figure S4. μBrain volume, atlas, cortical surfaces and microarray data
Figure S5: All ISH reconstructions.
Figure S6: PCA analysis of each brain specimen
Figure S7: Correlation between PC1 and age-related changes in expression in each tissue
Figure S8: Correlation between age-related gene expression changes across tissue
Figure S9: Developmental tissue enrichment of ZRT and non-ZRT genes
Figure S10: Gestational age distribution of fetal MRI scans (n=195) included in analysis.
Figure S11: Comparison of within- and across-subject estimates of cortical expansion.
Figure S12: Tissue and cell expression of $ZRT_{neo}$ genes at 15 PCW.
Figure S13: SATB2 ISH across cortical regions.
Figure S14: Chromatin accessibility near to ZRT genes in the fetal brain.
Figure S15: Expression of OPC markers in the intermediate zone.
Figure S16: OLIG1 expression in prenatal bulk tissue mRNA data.
Figure S17: image registration pipeline and template generation.
Figure S18: Cortical surface area scaling after removing repeated scans.
Figure S19: Proportion of vertex outliers in surface area data

**Supplemental references**

### Supplemental methods

#### GAN model architecture and training

We used the *pix2pix* architecture (**Figure 1b**) to synthesise Nissl-stained images from label annotations in order to replace artefacts within tissue sections.

##### *Model architecture*

The *pix2pix* model is a conditional generative adversarial network (GAN) trained on paired examples to perform image-to-image translation by combining a generator, $G$, with a discriminator, $D$. Following Isola et al.,[1] our generator network takes the form of a U-Net, with a symmetric encoder-decoder structure, and skip connections between corresponding encoding and decoding paths. The encoder was parameterised with 7 resolution levels, each comprising a 2D convolution [kernel size=(4,4), stride=(2,2); filters=(64, 128, 256, 512, 512, 512, 512)], batch normalisation and a leaky ReLu activation function (α=0.2). This is followed by a bottleneck layer composed from one 2D convolution with 512 filters and ReLu activation. The decoder layers consisted of 2D upsampling (implemented with nearest neighbour interpolation), followed by a 2D convolution [kernel size=(4,4), stride=(1,1), filters=(512, 512, 512, 512, 256, 128, 64)] with dropout (p=0.5) (**Figure 1b**). A final upsampling and convolution with *tanh* activation was applied to generate the 3-channel RGB output image.[58]

The *pix2pix* discriminator, *D*, is trained as a convolutional *PatchGAN.* By acting on small sections of an image rather than the whole image, this approach focuses on high-frequency image structure and texture while requiring fewer parameters.[1] In our application, the discriminator comprised three 2D convolutional layers [kernel = (4,4), stride = (2,2), (1,1) and (1,1), filters = 64, 128 and 1, respectively] with leaky ReLU (first two layers) and sigmoid (final layer) activations (**Figure 1b**). Batch normalization was applied after the second convolution. The final output is a 128 × 128 image with real/fake predictions for each 16 × 16 patch in the original input image.[1] We implemented the model in Python 3.7 using Tensorflow (2.4.1) and the Keras API [https://keras.io/].

##### *Training data*

We trained the *pix2pix* model on 1000 pairs of 256 × 256 image patches from 20μm resolution Nissl-stained sections and corresponding label annotations (**Figure 1a,b**). Each pair of training patches was visually inspected to ensure no tissue artefacts were present in the histological data and good alignment was observed between tissue section and anatomical labels. Pairs that failed visual inspection were rejected and replaced until 1000 pairs were selected. Pairs were automatically excluded if >75% pixels were labelled as background. Patches were randomly sampled from n=73 sections, with the remaining n=8 sections forming a validation set to evaluate model performance. Validation sections were spread evenly through the cerebral hemisphere. To increase the size and diversity of our training set and limit overfitting, we applied common data augmentation steps including random crop, jitter and flipping of images during training.

##### *Model training*

The model was trained in steps, alternating between training the generator and the discriminator with real and synthetic samples. As in Isola et al.[1] the discriminator loss was divided by 2 to slow down its learning rate compared to the generator and both *D* and *G* were trained using the Adam optimiser ($\beta_1 = 0.5$, $\beta_2 = 0.999$) with a learning rate of 0.0002 and batch size of 1 for a total 100 epochs. We set the regularisation parameter, λ, to 1. Alternative parameter settings are explored in **Figure S2**. Finally, the 3-channel RGB label images used for training were transformed to 21-

channel images (20 tissue labels + 1 background label) with each channel containing 1s in pixels belonging to a given label (0 otherwise).

##### *Model evaluation*

We retained eight full histological sections spread evenly through the cerebral hemisphere as a validation set. No image patches used for training were drawn from this dataset. To validate model performance, we split each Nissl-stained section and its corresponding anatomical label image into non-overlapping 256 ×256 patches. Labelled patches were used to generate new synthetic Nissl-stained patches by the trained generator.

To quantify visual and textural similarity between synthetic and ground truth Nissl images, we converted each image patch from RGB to HSV (Hue, Saturation, Value) format and calculated the similarity between hue and saturation values across both patches as:

$$\mathrm{sim}(F_1, F_2) = \frac{1}{1 + \sqrt{\sum (F_1 - F_2)^2}}$$

Where $F_1$ and $F_2$ are the vectors of pixel hue or saturation values for ground truth and synthetic images patches, respectively, normalised to unit length. As an additional measure of generator performance, we calculated a 'perceptual' similarity between ground truth and synthetic image patches based on high-level features of a large image recognition model pretrained on the ImageNet dataset (VGG19).[2–4] After removing the fully-connected classification layers of the pre-trained VGG19 model, each image patch (real and synthetic) was passed through the network, resulting in a 512 length vector output by the final layer to act as a high-level feature representation of each patch. Similarity between feature vectors was calculated as above, baseline measures of hue, saturation and perceptual similarity were calculated after randomising pixels within each ground truth patch.

#### Template construction

Following automated repair of major tissue artefacts present in the data, we aimed to develop a 3-dimensional reconstruction of the fetal brain to facilitate comparison with *in vivo* MR imaging data.

##### *Slice-to-slice alignment*

Repaired Nissl-stained images were converted to greyscale and padded with a 200 pixel zero-filled border to allow for large translations while retaining the section in the field-of-view during image registration. Corresponding label images were converted from RGB format to one-channel label images, with labels numbered according to a look-up table (**Table S2**).

For initial alignment, we implemented a graph-based, slice-to-slice registration to a chosen reference section via shortest-path transforms.[5,6] The central section was chosen as the reference, and pairwise rigid transforms were estimated between each section and up to five neighbouring sections in the direction of the reference. After each transform, a weight is calculated between the aligned sections:

$$\omega_{i,j} = (1 - r_{i,j}) \times (1 + \lambda)^{|I_i - I_j|}$$

Where the weighting, ω, between two aligned images, $i$ and $j$, depends upon their pixelwise correlation, $r$, after alignment weighted by the number of intermediate sections between them (based on the absolute difference of section index, $I$). The hyperparameter, $\lambda$, was set to 0.5 and acts as a penalty on skipping slices, with a higher value penalising transforms that skip intermediate sections.[5,6] Dijkstra's shortest-path algorithm was used to calculate the set of transforms with lowest cost to align a given section to the reference.[5,6] The selected transforms were then composed and applied to both the image and its corresponding labels to bring all sections into approximate alignment with the central slice.

Aligned sections were stacked along the anterior-posterior axis into a 3D array with resolution 0.02 × 0.02 × 0.5mm. Section thickness was determined by the sampling strategy of the original data (detailed in Ding et al.[7]). Consecutive sections included in the atlas were spaced approximately 0.5 mm apart (**Table S1**) and were thus assigned a nominal thickness of 500$\mu$m. Due to the sampling strategy employed during tissue sampling and histology, tissue sections were sparsely sampled along the length of the cerebral hemisphere. Where full sections were missing, or excluded, or the distance between adjacent slices was larger than 0.5mm, we repeated the preceding sections were up to 5 times to fill the gap. This strategy was selected to limit abrupt transitions between neighbouring sections, preserve the overall volume of the hemisphere and allow for volumetric registration in subsequent steps. To account for missing slices at the anterior and posterior poles, manual labels were drawn following the tissue contours of the adjacent slices to create a synthetic cortical label. The trained *pix2pix* generative model was then used to generate a Nissl-stained coronal section to append to each pole. This process resulted in a 3D NIFTI volume of voxel size 1420 × 2678 × 125 and voxel dimensions 0.02 × 0.02 × 0.5mm (**Figure 1e; Figure S17**).

*Affine registration to a fetal brain shape reference*

We employed a population-based average anatomical image: specifically the 22-week timepoint of the Gholipour et al. spatio-temporal fetal MRI atlas as a shape prior for 3D reconstruction (**Figure S3**).[8]

The Gholipour MRI atlas contained T2-weighted anatomical fetal MRI templates and a set of 50 anatomical brain tissue labels at 1mm isotropic voxel resolution. We downloaded the T2-weighted template image and accompanying tissue labels for the 22-week timepoint, removing extracerebral CSF, midbrain and cerebellar structures and matching tissue labels of the MRI atlas to the µBrain tissue labels. We upsampled the MRI template and label images to 50µm isotropic resolution before cropping and rotating into approximate alignment with the 3D µBrain volume. We converted the MRI volume into a Nissl-like contrast using the trained GAN model by slicing the label image coronally and passing each slice as input to the trained model (**Figure S3c-d**). Nissl-contrast images were re-stacked into a 3D volume and used as an anatomical prior for registration.

Using the shape prior, we performed an iterative affine registration procedure[5,6] by first estimating an affine alignment of the MRI-based anatomical prior to the histological volume. The transformed anatomical prior was then resliced, with each coronal section acting as a target for 2D registration with the corresponding histological section. The registered histological sections then form the target for the next 3D registration (**Figure S17**). This process was repeated for a total 5 iterations, producing a final 3D volume with aligned coronal slices and a global shape approximately matched to the *in utero* fetal brain (**Figure 1e**). A final 2D affine registration was calculated between the original and final aligned histological sections, and the Nissl-contrast images and corresponding tissue labels transformed into the 3D volume.

*Final template construction*

Typically, brain templates are probabilistic estimates constructed from multiple individual datasets, representing a population-average anatomy, the principal benefit of which is to provide a common coordinate space for analysis and remove bias towards any individual's brain anatomy.[9] Borrowing from this philosophy, we framed the output of the preceding steps: a 3D volume with approximately aligned coronal sections and global shape, as a single possible representation of the ground truth cerebral volume that captures various idiosyncrasies of the reconstruction pipeline including tissue sectioning frequency and selection, image repair, registration and/or potential misalignment. We employed a data augmentation technique to create multiple instances of the volume reconstruction, created nonlinear deformations by deleting and/or repeating up to 25 randomly selected slices along all 3 image dimensions and forming a population of n=50 unique representations of the original data. Matched augmentations were applied to both the histological reconstruction and accompanying anatomical labels (**Figure S17**).

For each augmented volume, we resampled the in-plane, coronal resolution to $150 \times 150\mu$m resolution (slice thickness = $500\mu$m) and, for each coronal slice, performed a weighted nonlinear registration to neighbouring sections (symmetric normalisation [SyN] metric=cross-correlation; flow_sigma=3.0, total_sigma=1.0, grad_step=0.25). Adjacent sections (up to 8 neighbours) were weighted based on distance to the source section. After registration, the halfway transform was applied to each section. Section-to-section registration and transformation was performed once in the anterior-posterior direction, before repeating in the posterior-anterior direction. For each volume, slice-to-slice nonlinear registrations were calculated for a total of 3 iterations.

Finally, to create a smooth 3D reconstructed volume, we co-registered all 50 augmented, aligned volumes into a single probabilistic anatomical template with voxel resolution $150 \times 150 \times\ 150\mu$m using iterative, whole-brain nonlinear registration (SyN metric=cross-correlation; iterations=3; grad_step=0.25) (**Figure S17**). Transforms were applied to each of the corresponding anatomical label volumes and majority vote used to create a final set of brain tissue and cortical labels (**Figure S4**). All image registration was performed in Python 3.7 using *antspyx* (0.2.7).[10]

##### *Cortical reconstruction*

To reconstruct the fetal cortical surface, we adapted existing protocols for *ex vivo* [https://freesurfer.net/fswiki/ExVivo] and non-human primate [https://prime-re.github.io/] surface reconstruction with Freesurfer.[11] We first created a 'dummy' volume with 1mm voxel dimensions and created a 'white matter' mask by combining tissue labels for all subcortical structures and tissue zones excluding the cortical plate. This volume was tessellated (Freesurfer commands: *mri_pretess*, *mri_tesselate*), and the initial surface smoothed (*mris_smooth*) and inflated to a sphere (*mris_inflate*, *mris_sphere*). Topological errors in the initial surface estimates were detected and fixed using manual edits to the brain and white matter masks, before repeating the process. Finally, a pseudo-T2 volume was created using tissue labels, assigning all voxels in the cortical plate intensities expected in grey matter by Freesurfer. This volume was used to generate inner and outer surfaces for the volume (*mris_make_surfaces*). Surfaces were smoothed for 50 iterations and inspected for topological errors, before conversion to *gifti* format and rescaling to the original size (**Figure 1f, S4**). All processing was performed with Freesurfer (7.3.2).

### Supplemental figures

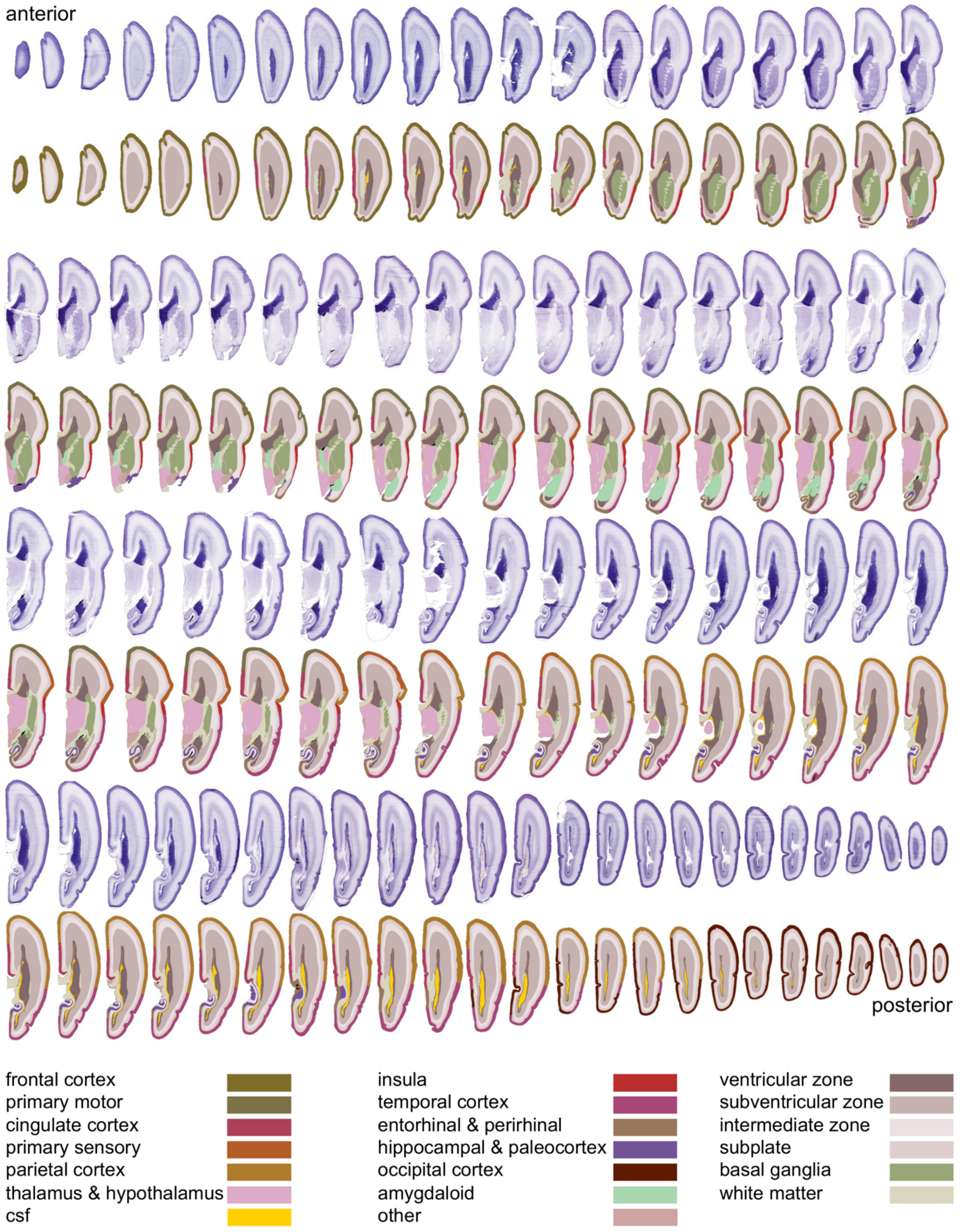


**Figure S1: Lightbox of reference atlas sections**

Serial Nissl-stained sections and corresponding anatomical annotations used to construct the $\mu$Brain atlas. Annotations represent a simplified label set (brain-labels) based on the hierarchical ontology of the BrainSpan reference atlas (**Table S2**).

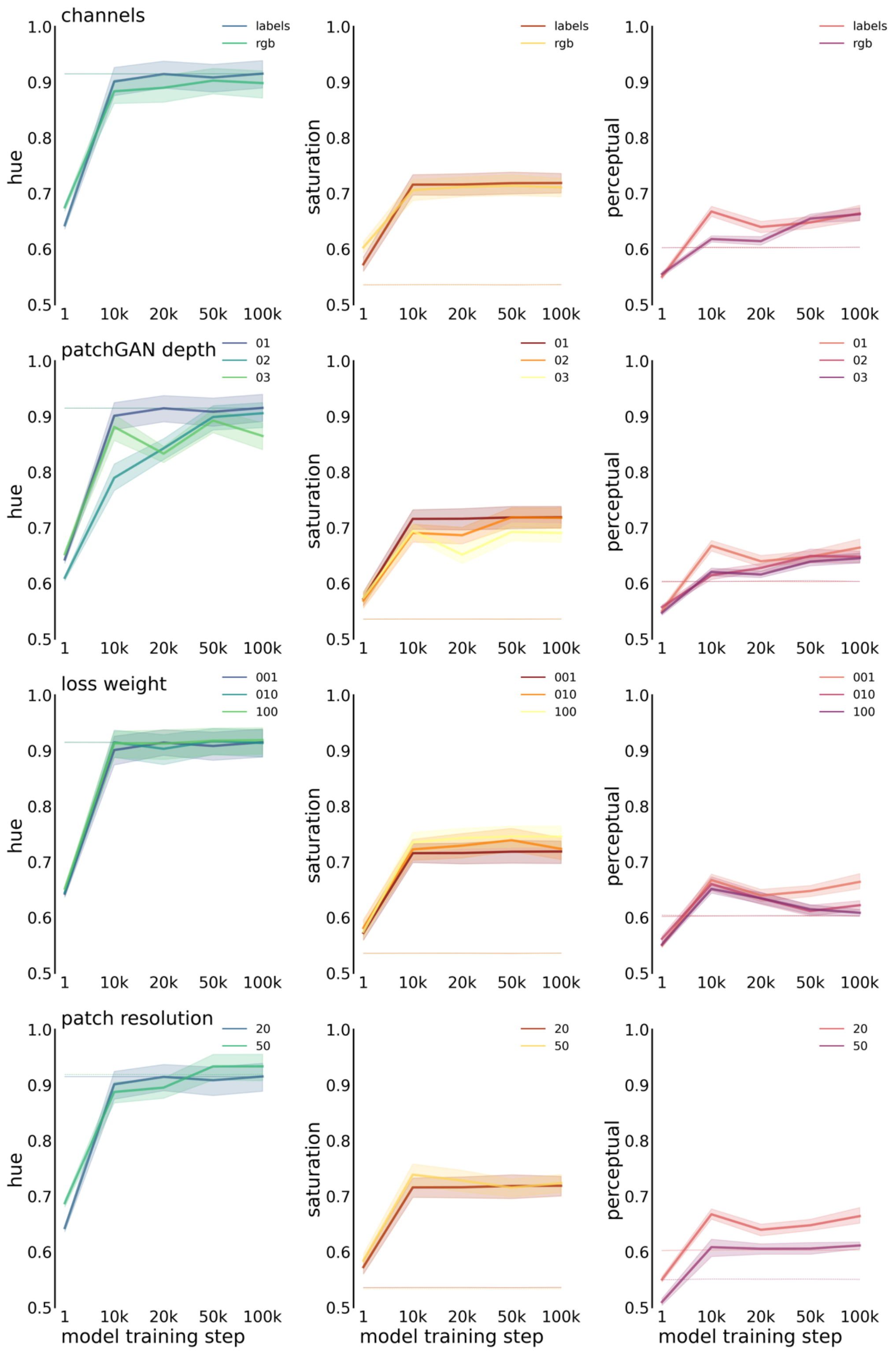


**Figure S2: Model performance over hyperparameters**

Mean similarity of hue (left) and saturation (middle) of patch predictions compared to ground truth during model training with different parameters (patch resolution in $\mu$m; loss weight; number of *patchGAN* layers and input channels). Perceptual similarity (right) was calculated based on the similarity of outputs from a pretrained VGG19 model. Baseline measures of similarity were generated by randomising pixels within each patch (dashed lines).

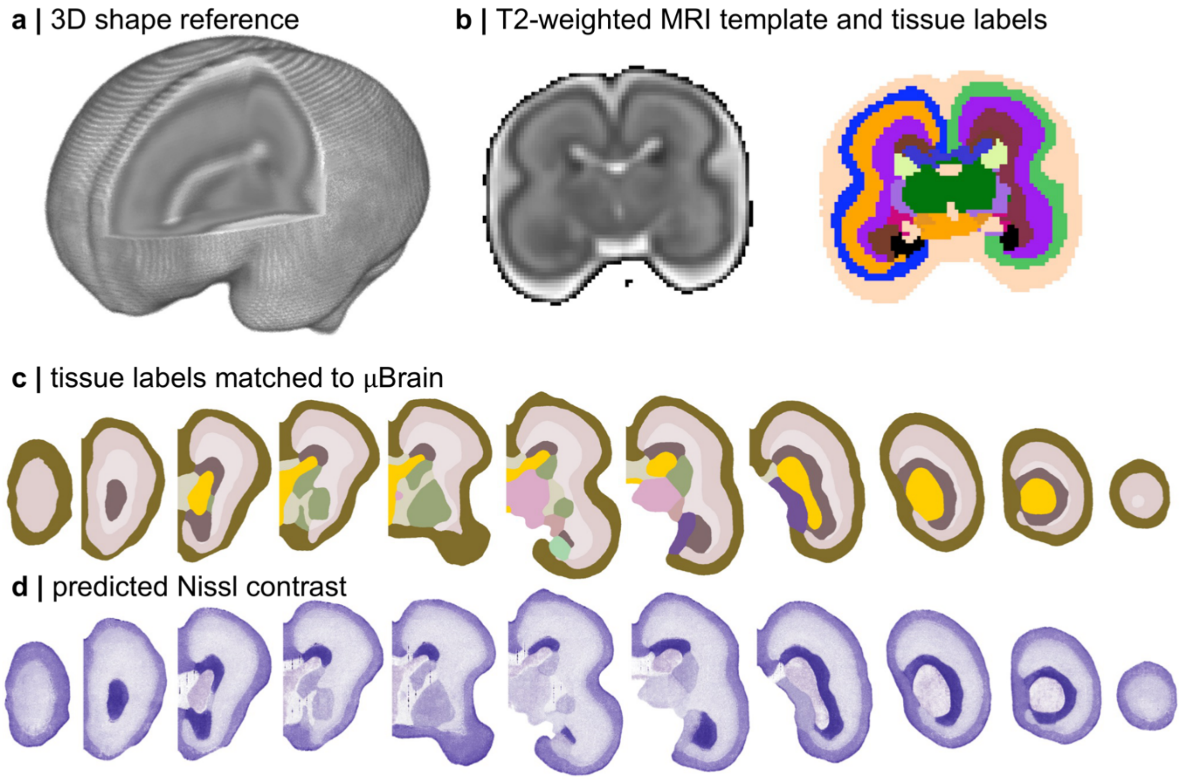


**Figure S3: Shape reference for affine registration**

a. A fetal brain atlas at 22 gestational weeks was used as a shape reference for affine registration. b. manual tissue labels were matched to $\mu$Brain anatomical labels (c) and used to generate synthetic 'Nissl-contrast' sections (d) to act as a target for registration.

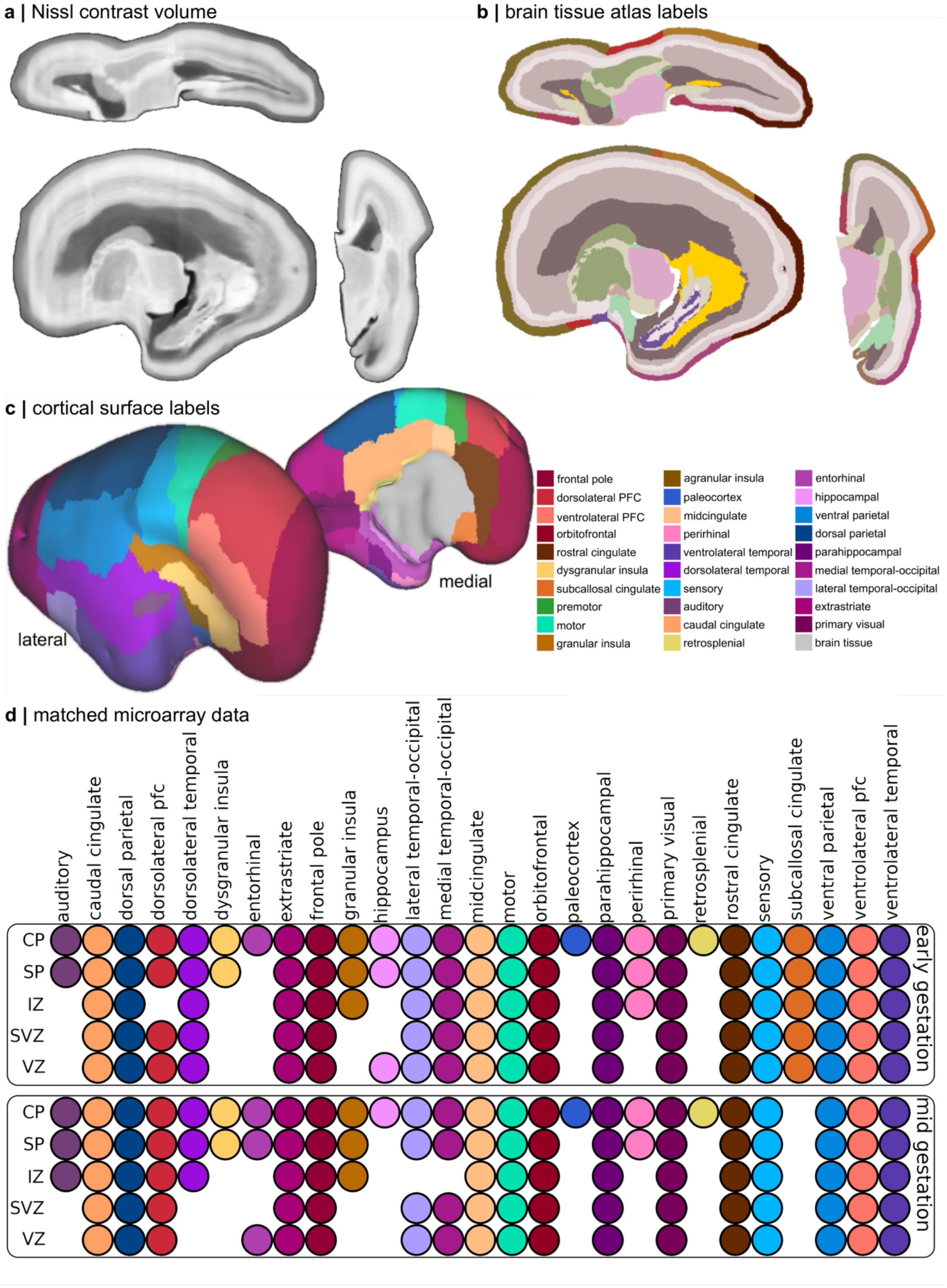


**Figure S4. $\mu$Brain volume, atlas, cortical surfaces and microarray data**

a. 3D volumetric reconstruction of histological data. b.corresponding anatomical atlas labels (brain-labels; **Table S2**). Coloured as in **Figure S1**. c. lateral and medial cortical surface reconstruction labelled by cortical area (cortex-labels; **Table S2**). d. available LMD microarray data matched to cortical labels at early and mid gestational timepoints (15/16PCW and 21PCW, respectively) for each of five tissue zones. CP: cortical plate; SP: subplate; IZ, intermediate zone; SVZ: subventricular zone; VZ: ventricular zone

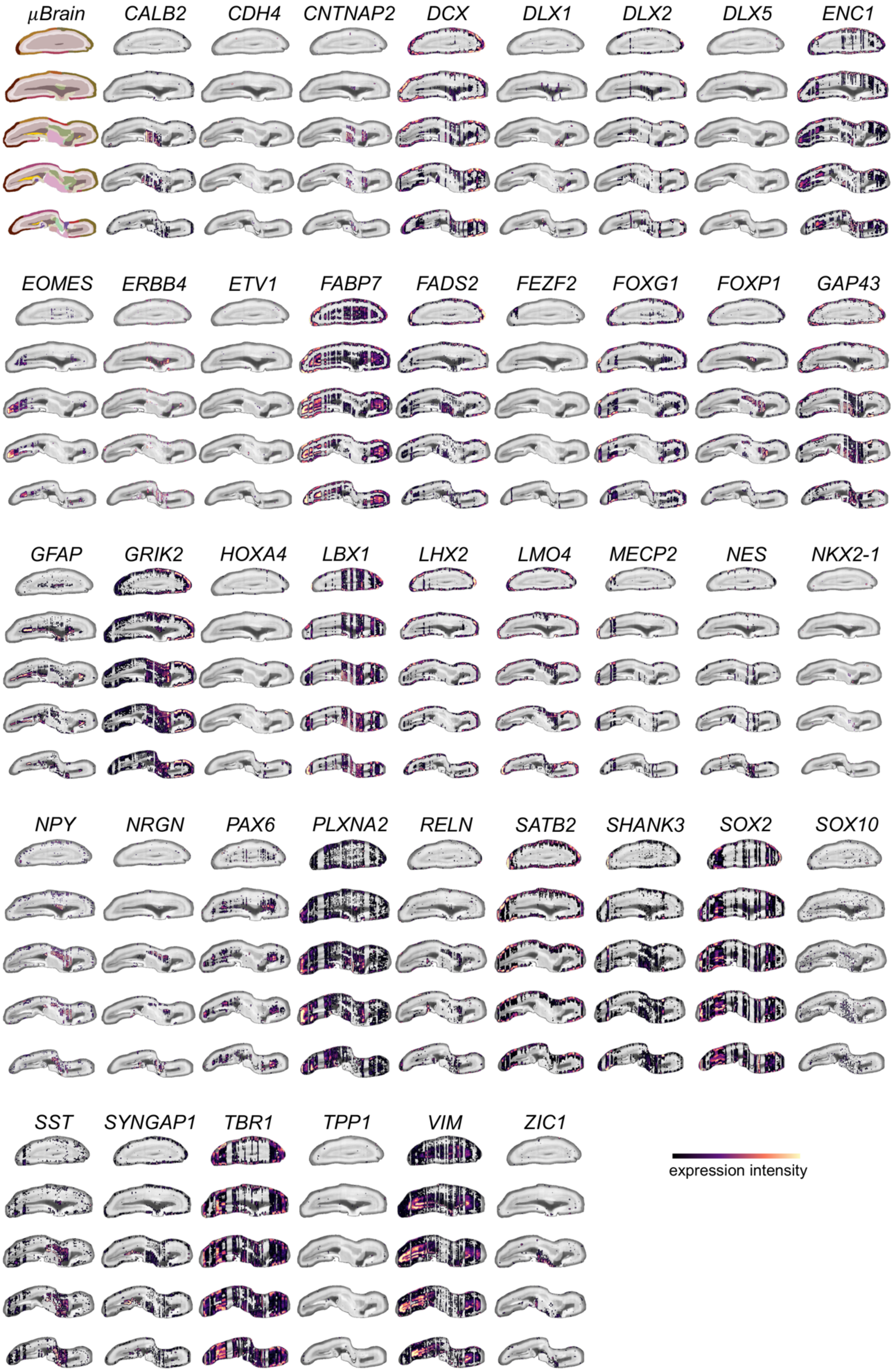


**Figure S5: All ISH reconstructions.**

Axial slices from partial 3D reconstruction of ISH expression data from n=41 neurodevelopmental genes.

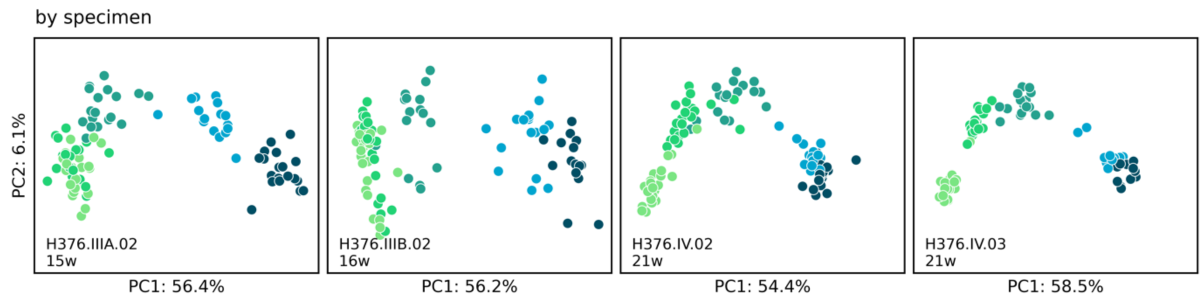


**Figure S6: PCA analysis of each brain specimen**

PCA was performed separately on each specimen's micrparray data. The first two components are shown for each specimen.

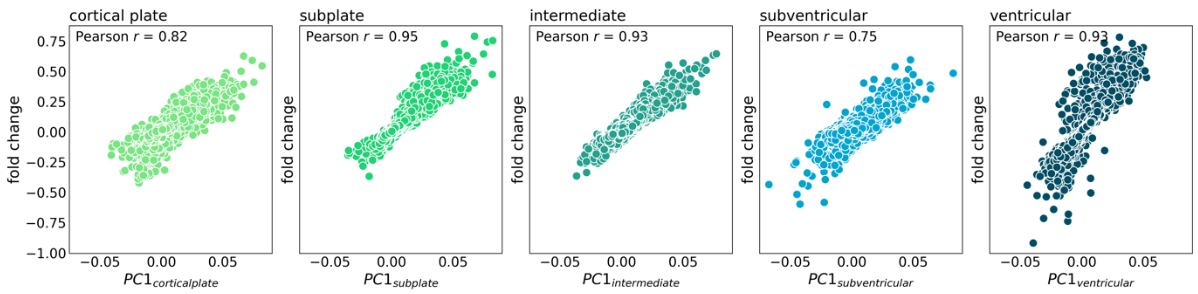


**Figure S7: Correlation between PC1 and age-related changes in expression in each tissue**

For each tissue zone, we calculated the correlation between age-related changes (between 15 and 21PCW) and the first principal component. Age related change explained the majority of variance in each tissue zone.

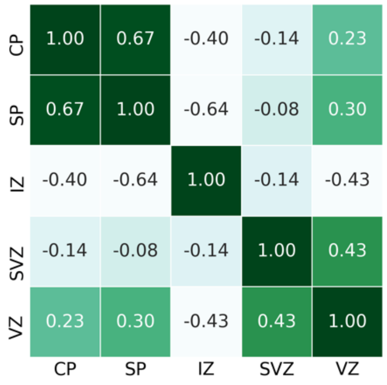


**Figure S8: Correlation between age-related gene expression changes across tissue**

Correlation between age-related changes in each gene across tissue zones. Age-related changes in gene expression were most similar between neighbouring tissue zones.

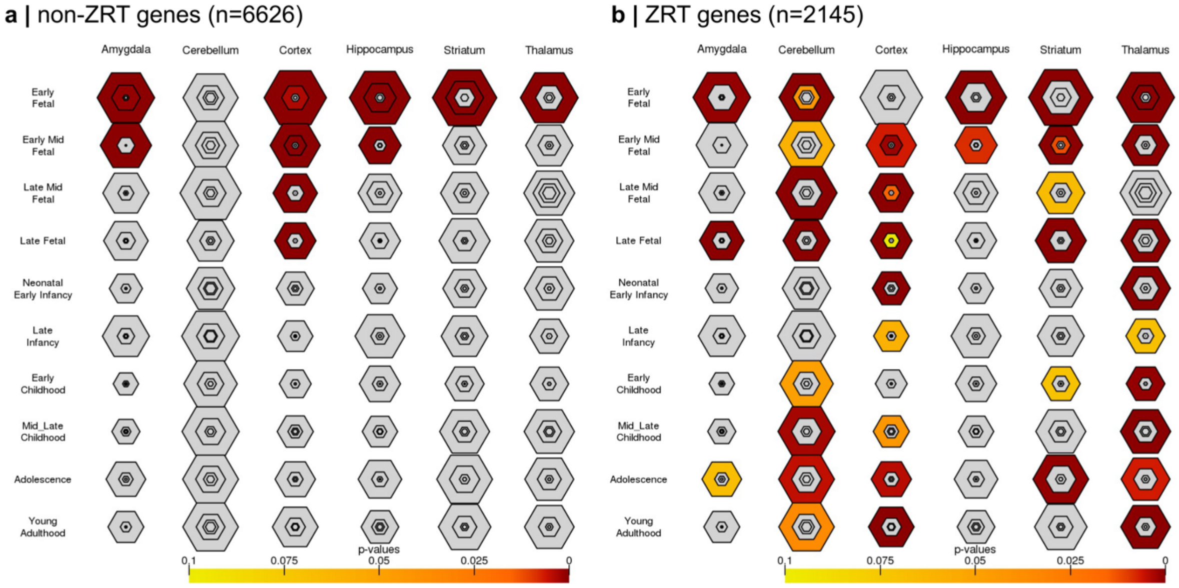


**Figure S9: Developmental tissue enrichment of ZRT and non-ZRT genes**

Output from Cell-Specific Enrichment Analysis[12] of ZRT and non-ZRT genes. Significant, cell-specific enrichment of genes are illustrated by hexagon size and colour.

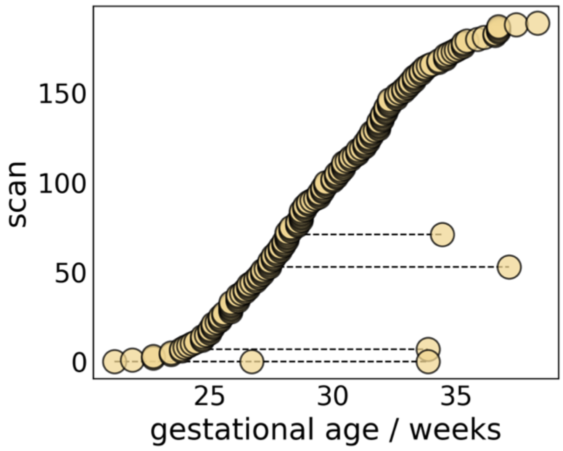


**Figure S10: Gestational age distribution of fetal MRI scans (n=195) included in analysis.**

Four fetuses were scanned more than once during gestation

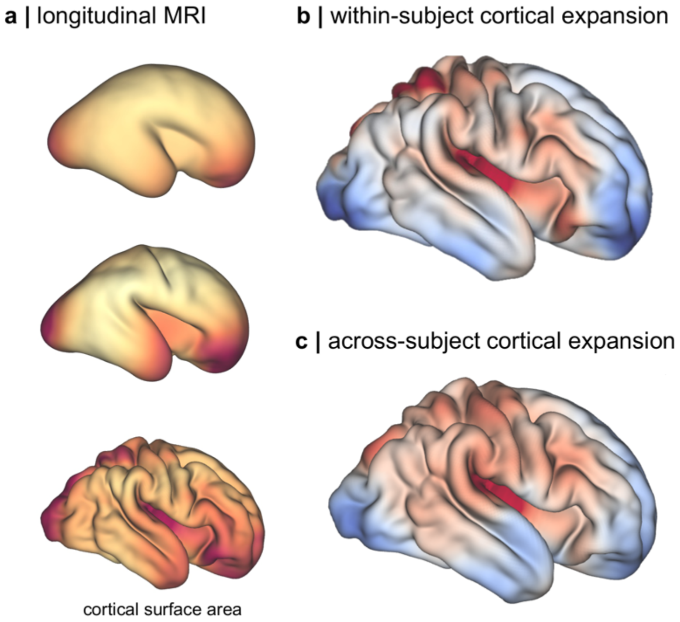


**Figure S11: Comparison of within- and across-subject estimates of cortical expansion.**

a. A single fetus was scanned three times during gestation After MRI processing, cortical surfaces were extracted from each scan and vertex area was calculated after resampling native topology onto the dHCP template surface. b. Differences in proportional area, calculated as the area at each vertex divided by total cortical surface area, between the earliest and latest timepoints. Warm colours indicate a preferential increase in regional surface area over time. c. Whole-group, cross-sectional expansion map from **Figure 3**.

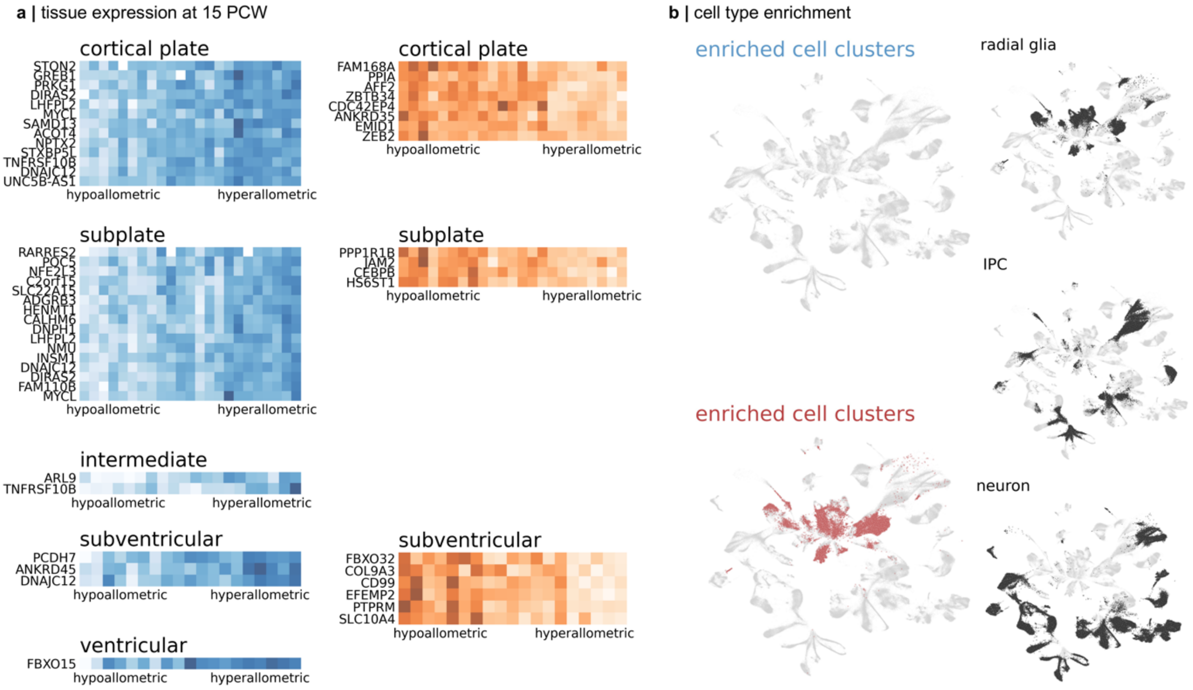


**Figure S12: Tissue and cell expression of ZRT$_{neo}$ genes at 15 PCW.**

**a.** normalised (Z-score) expression profiles for genes correlated with areal scaling in each tissue zone at 15PCW. Negative associations (higher relative expression in hypoallometric regions) shown in blue, positive associations are in red. Lighter colours indicate higher relative expression. **b.** mid-gestation cell clusters[13] significantly enriched for genes associated with areal scaling in at 15PCW. Territories of three canonical cell types are shown.

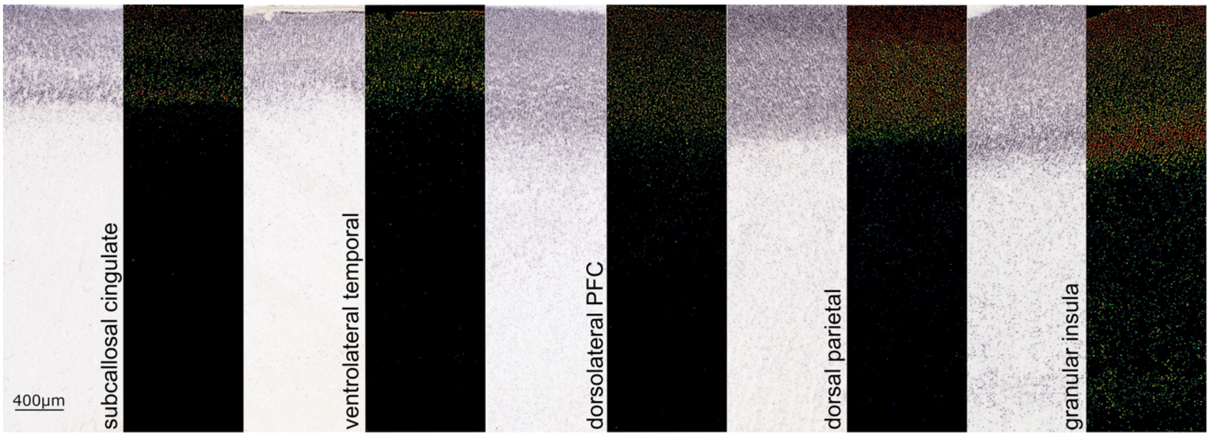


**Figure S13: SATB2 ISH across cortical regions.**

Example ISH staining of SATB2+ cells across the cortical anlage in five cortical regions.

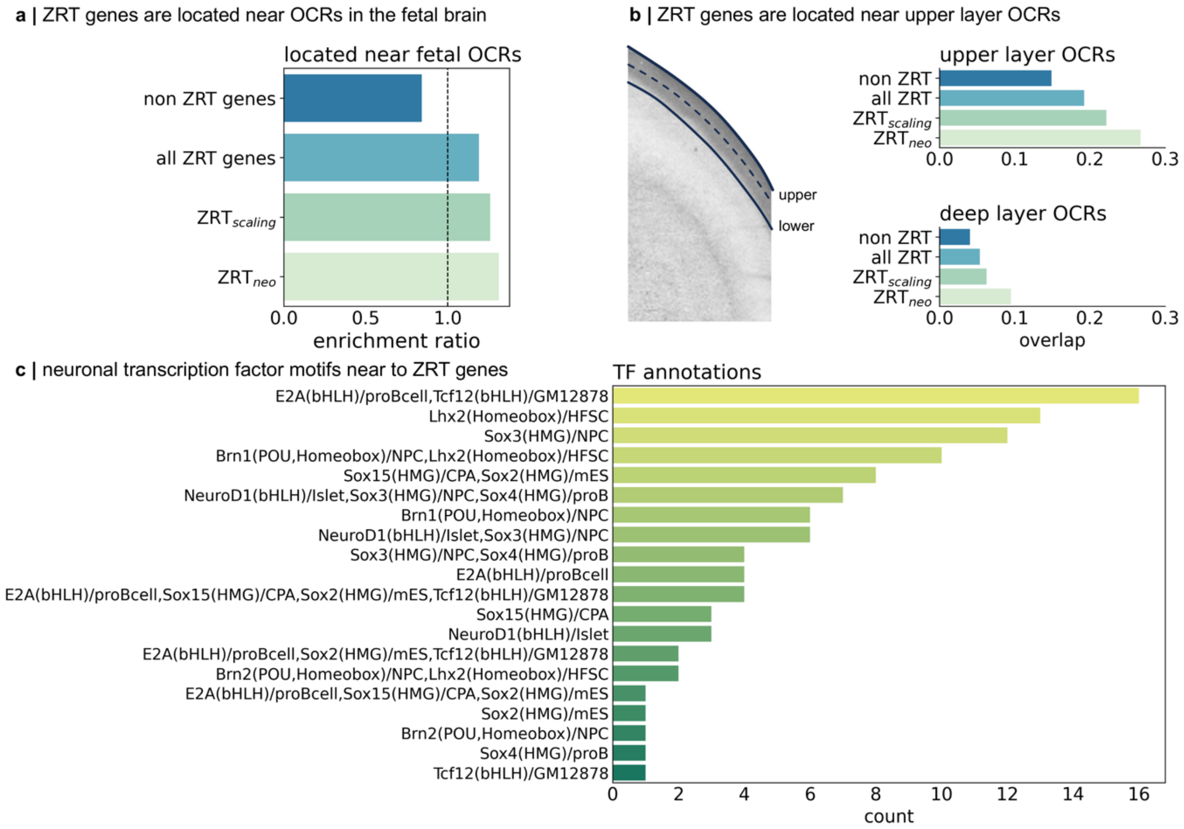


**Figure S14: Chromatin accessibility near to ZRT genes in the fetal brain.**

**a**. enrichment of different ZRT genesets in all genes located near to OCRs in the prenatal brain. **b.** enrichment in laminar-specific OCRs. **c.** number of transcription factor motif annotations located near to ZRT genes.

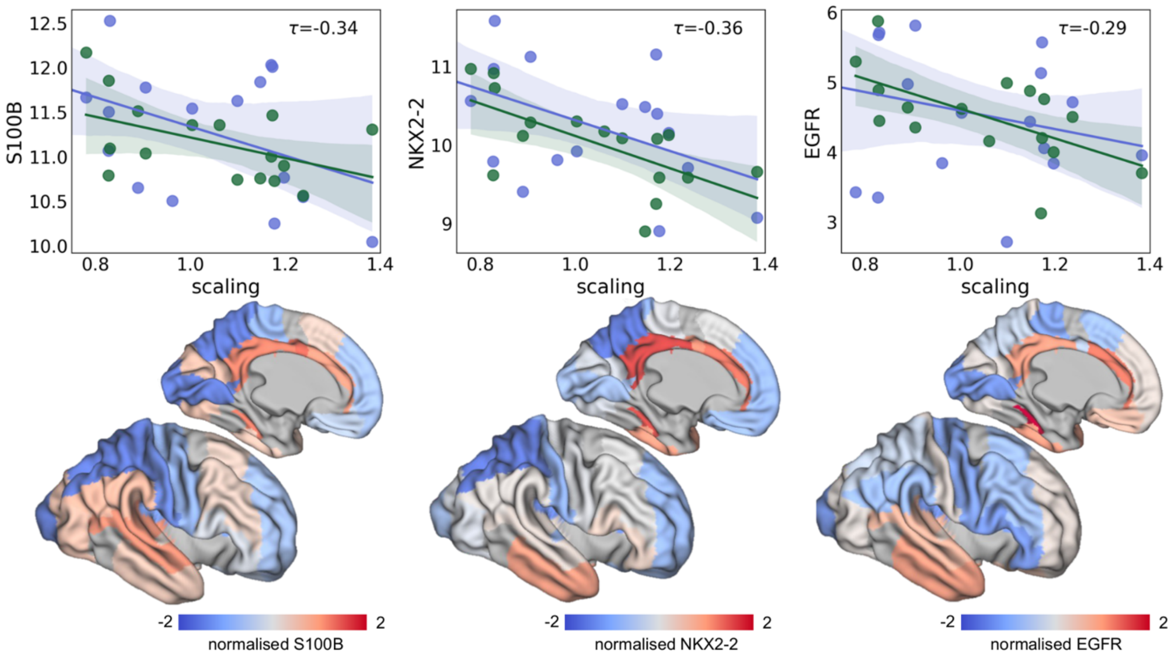


**Figure S15: Expression of OPC markers in the intermediate zone.**

Top: regional expression of oligodendrocyte-lineage markers (S100B, NKX2-2, EGFR) in the intermediate zone of two mid-gestation brains is associated with cortical expansion (scaling coefficient). Bottom: Mean normalised expression of each marker in the IZ calculated within each $\mu$Brain label and projected onto the 35w surface template.

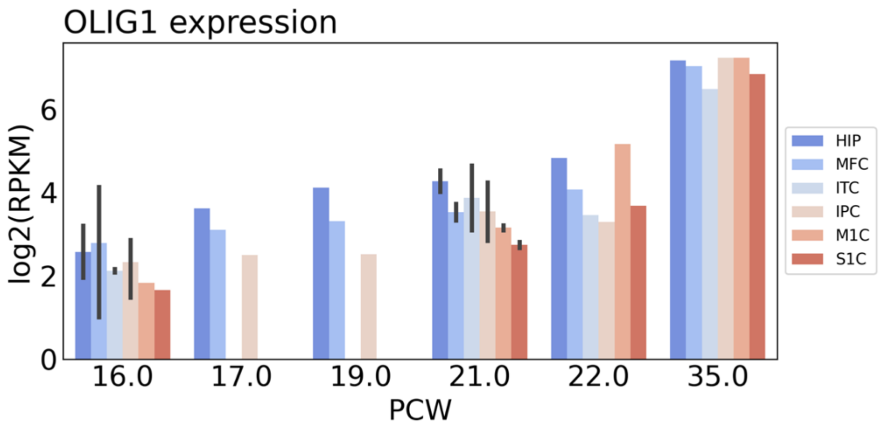


**Figure S16: OLIG1 expression in prenatal bulk tissue mRNA data.**

Regional estimates of OLIG1 expression were extracted from six cortical regions with differential allometric expansion (hypoallometric=blue; hyperallometric=red) for n=9 prenatal brain specimens aged 16 – 35PCW. HIP: hippocampus; MFC: medial frontal cortex; ITC: inferior temporal cortex; IPC: inferior parietal cortex; M1C: primary motor cortex; S1C: primary sensory cortex.

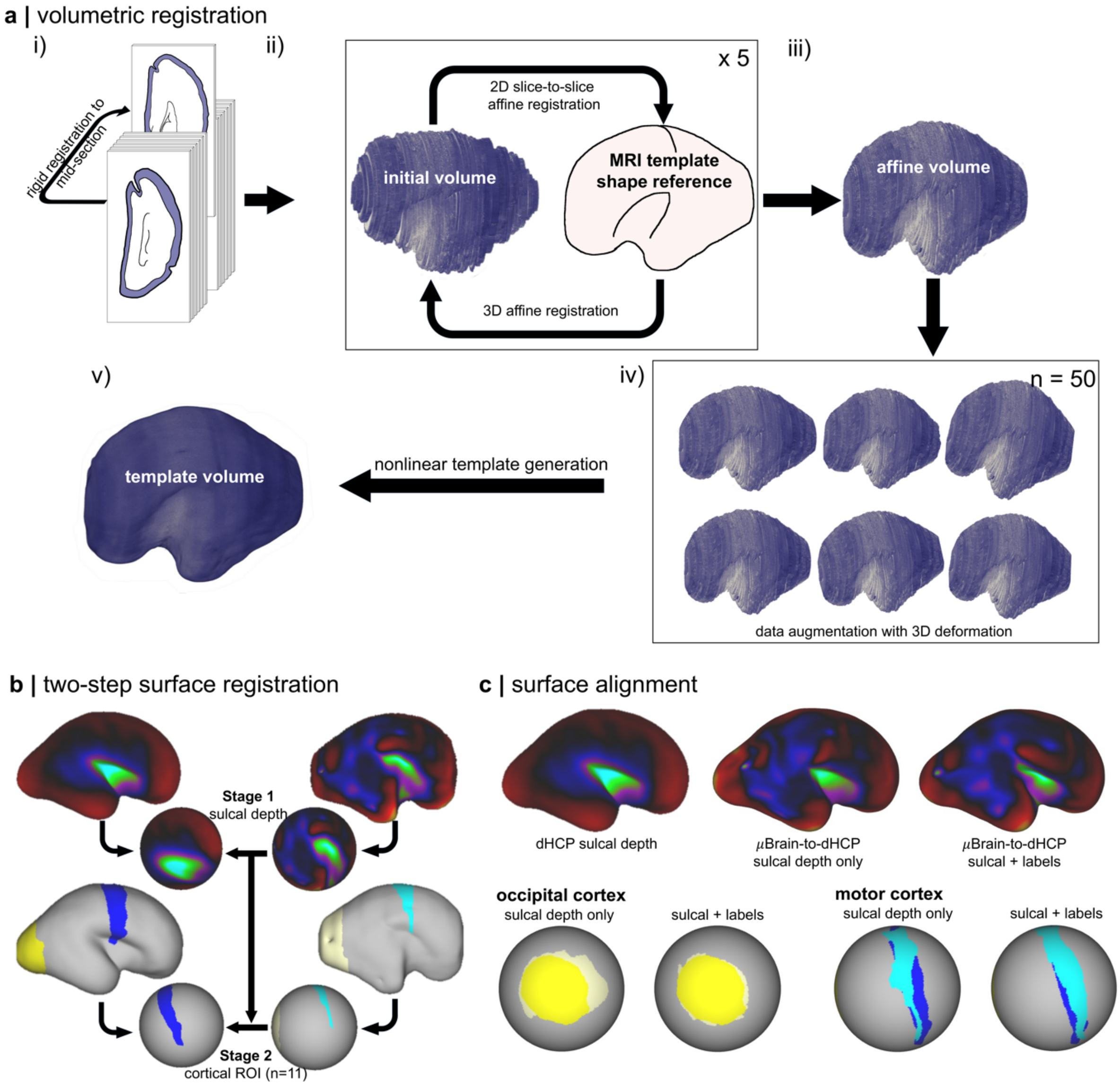


**Figure S17: image registration pipeline and template generation.**

**a**. clockwise: i) Repaired Nissl sections were first aligned to the mid-section using a graph-based rigid registration algorithm and stacked into a 3D volume. ii) An MRI-based anatomical template was used as a shape reference to guide affine alignment between sections. After transforming each slice of the template to a Nissl-like contrast using the trained *pix2pix* model (**Figure S3**), we aligned the reference to the Nissl volume using a 3D affine registration. The aligned volume was resliced and 2D section-to-slice registration to the shape reference performed for each Nissl section. This process was repeated for 5 iterations to form an 'affine volume' (iii). iv)We employed a data augmentation to induce 3D deformations in the affine volume and create a population (n=50) of alternative representations of the cerebral volume, resampled to 150$\mu$m resolution. For each volume, each section was registered to its neighbours using nonlinear registration. Registrations were performed slice-by-slice, once forward and once backwards along the volume for a total of 3 iterations. v) All 50 volumes were co-registered and averaged to create a final 3D template using symmetric nonlinear registration. **b.** A two-step registration process was used to align the $\mu$Brain cortical surface to the dHCP fetal template. Nonlinear surface registrations were performed between spherical representations of each surface using MSM driven by sulcal depth (Stage 1). This alignment was used to initialise a second registration driven by a set of matched cortical regions-of-interest (n=11; Stage 2). Only the occipital (yellow) and motor (blue) regions are shown for simplicity. Each cortical region was passed to the algorithm as a separate metric with alignment jointly optimised over all regions. **c.** cortical surface alignment after Stage 1 (sulcal depth only) and Stage 2 (sulcal depth +

cortical labels). Top row shows sulcal depth maps on the fetal surface. Bottom rows shows alignment of occipital and motor regions on the spherical surface.

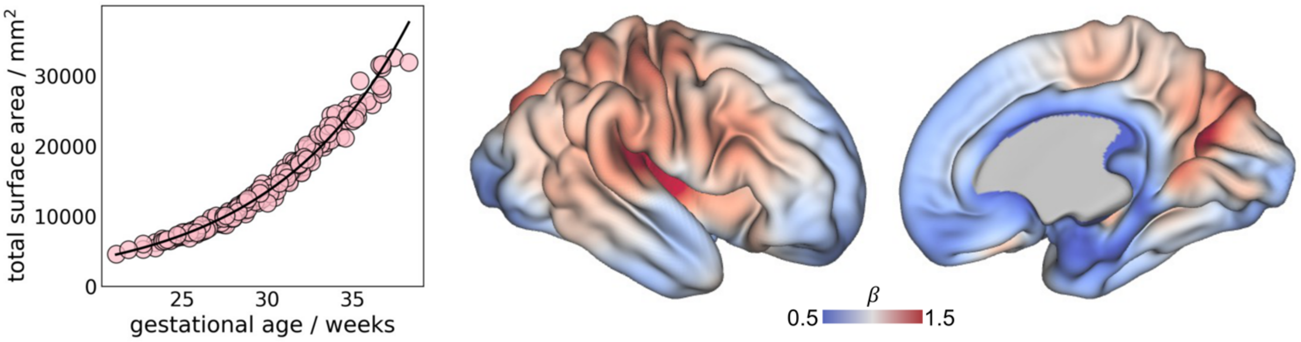


**Figure S18: Cortical surface area scaling after removing repeated scans.**

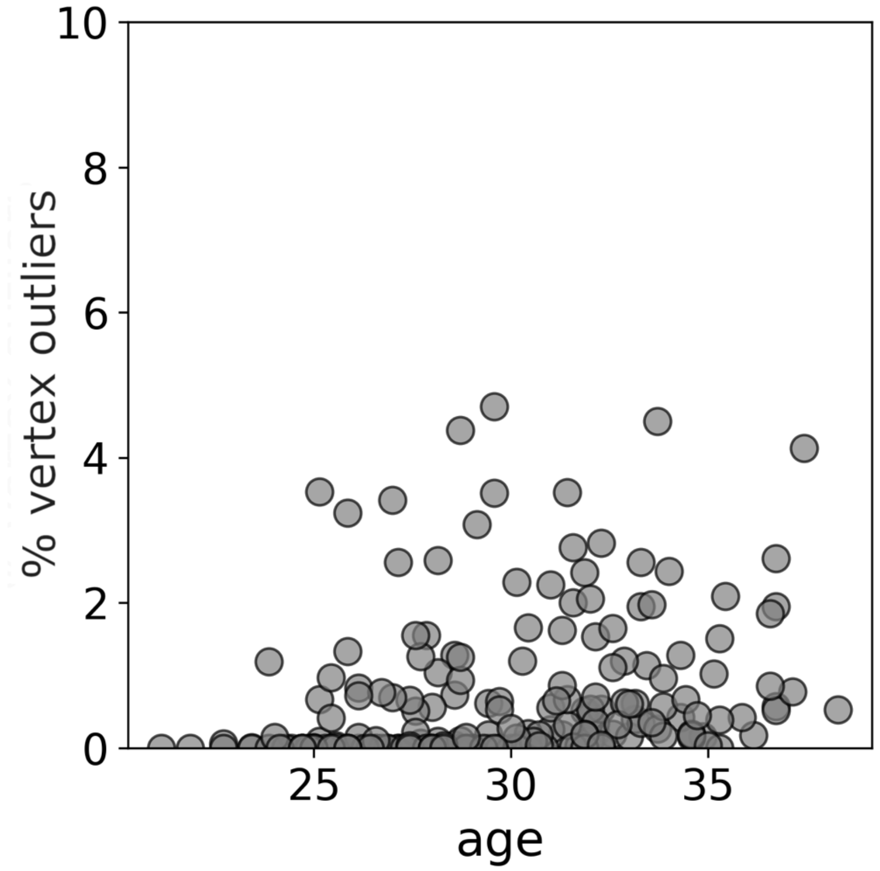


**Figure S19: Proportion of vertex outliers in surface area data**
